# Greenhouse spatial effects detected in the barley (*Hordeum vulgare* L.) epigenome underlie stochasticity of DNA methylation

**DOI:** 10.1101/2020.04.20.051326

**Authors:** Moumouni Konate, Michael J. Wilkinson, Julian Taylor, Eileen S. Scott, Bettina Berger, Carlos Marcelino Rodriguez Lopez

**Affiliations:** Institut de l’Environnement et de Recherche Agricole (INERA), DRREA-Ouest, 01 BP 910 Bobo Dioulasso 01, Burkina Faso; Institute of Biological, Environmental and Rural Sciences, Penglais Campus, Aberystwyth, Ceredigion, SY23 3EB, UK; Biometry Hub, School of Agriculture, Food and Wine, Waite Research Institute, The University of Adelaide, Waite Campus, PMB 1, Glen Osmond, SA 5064, Australia; School of Agriculture, Food and Wine, Waite Research Institute, The University of Adelaide, Waite Campus, PMB 1, Glen Osmond, SA 5064, Australia; The Plant Accelerator, Australian Plant Phenomics Facility, School of Agriculture, Food and Wine, Waite Research Institute, The University of Adelaide, Waite Campus, PMB 1, Glen Osmond, SA 5064, Australia; Environmental Epigenetics and Genetics Group; Department of Horticulture, College of Agriculture, Food and Environment, University of Kentucky, Lexington, KY 40546, USA

**Keywords:** epigenetics, positional effect, phenotypic plasticity, genome by environment, salt stress, MSAP

## Abstract

Environmental cues are known to alter the methylation profile of genomic DNA, and thereby change the expression of some genes. A proportion of such modifications may become adaptive by adjusting expression of stress response genes but others been shown to be highly stochastic, even under controlled conditions. The influence of environmental flux on plants adds an additional layer of complexity that has potential to confound attempts to interpret interactions between environment, methylome and plant form. We therefore adopt a positional and longitudinal approach to study progressive changes to barley DNA methylation patterns in response to salt exposure during development under greenhouse conditions. Methylation-Sensitive Amplified Polymorphism (MSAP) and phenotypic analyses of nine diverse barley varieties were grown in a randomized plot design, under two salt treatments (0 mM and 75 mM NaCl). Combining environmental, phenotypic and epigenetic data analyses, we show that at least part of the epigenetic variability, previously described as stochastic, is linked to environmental micro-variations during plant growth. Additionally, we show that differences in methylation increase with time of exposure to micro-variations in environment. We propose that subsequent epigenetic studies take into account microclimate-induced epigenetic variability.

## 1 Introduction

Plant epigenetic mechanisms that can alter gene expression include the actions of short-interfering RNAs (siRNAs), chemical modification of histone tails and DNA methylation (Vanyushin, 2006; Sawan *et al*., 2008). These have been variously implicated in orchestrating developmental processes (Kohler and Makarevich, 2006; Ishida *et al*., 2008; Ay *et al*., 2014; Jung *et al*., 2015; Kooke *et al*., 2015), cell and organ differentiation (Joyce *et al*., 2003; Kohler and Makarevich, 2006; Kitimu *et al*., 2015; Kooke *et al*., 2015; Konate *et al*., 2020), reproduction (Yaish *et al*., 2011; Podio *et al*., 2014), parental imprinting (Gehring *et al*., 2006), acquired transgenerational trait inheritance (Tricker *et al*., 2013a; Tricker *et al*., 2013b) and adaptation to stress (Bird and Jaenisch, 2003; Boyko and Kovalchuk, 2008; Tricker *et al*., 2012).

DNA methylation has emerged as the prominent epigenetic signature for past or contemporary exposure of a plant to environmental insults (e.g. Xie *et al*. (2017) and has been implicated in the moderation of stress response (Bird and Jaenisch, 2003; Zilberman and Henikoff, 2007; Boyko and Kovalchuk, 2008). For instance, Tricker *et al*. (2012) reported that *Arabidopsis thaliana* responded to high relative humidity stress by suppressing the expression of two genes that control stomatal development through DNA methylation. DNA methylation has been similarly implicated in the response of various plant species to a range of stresses, including excess salt (Karan *et al*., 2012; Konate *et al*., 2018), temperature extremes (Steward *et al*., 2002; Bastow *et al*., 2004; Hashida *et al*., 2006; Pecinka *et al*., 2010; Song *et al*., 2012), herbivory (Herrera and Bazaga, 2011; Herrera and Bazaga, 2013) and heterogeneous environmental pressure (Wang *et al*., 2016). However, the relationship between DNA methylation and the stress effect is imprecise. Many of the methylation changes observed under stress fail to occur consistently across all genotypes or populations studied, and many others are not obviously associated with exonic regions. Fewer still can be directly tied to a particular stress response gene. Such observations have been described as stochastic (Karan *et al*., 2012; Tricker *et al*., 2012), spontaneous (Raj and van Oudenaarden, 2008; Becker *et al*., 2011; van der Graaf *et al*., 2015), and without clear triggering factors (i.e. occurring randomly in the genome independently of stress). Many have considered the random and spontaneous alteration of DNA methylation is an adaptive biological process in its own right; one that drives diversity and evolution in a Lamarckian-like fashion (Feinberg and Irizarry, 2010; Meyer and Roeder, 2014; Soen *et al*., 2015; van der Graaf *et al*., 2015; Vogt, 2015) and with the clear potential to alter fitness (Consuegra and Rodríguez López, 2016). Additionally, Soen *et al*. (2015) proposed a conceptual framework of random variations in the genome, instigated in response to environmental cues. They hypothesized that imposition of diverse types of stress upon individual organisms during development gives rise to an adaptive improvisation which deploys random phenotypic variations that allows some individuals to cope with unstable ambient conditions. However, the authors did not suggest an epigenetic mechanism that might be involved in the regulation of such adaptive phenotypic variation.

In a pivotal piece, Vogt (2015) provided insight into the concept of random variability. The author linked ‘stochastic developmental variation’ to stochastic occurrence of DNA methylation (Bird and Jaenisch, 2003; Field and Blackman, 2003). However, Vogt did not consider in depth the possible role that microclimatic variation may play in this apparent stochasticity. Herrera and Bazaga (2010) suspected a role for mesoclimate in driving the epigenetic variability of natural populations but did not anticipate marked environmental differences to occur under controlled experimental conditions (greenhouse, growth room).

Moreover, since genome-by-environment interactions have been shown to be at least partially regulated by DNA methylation (Verhoeven *et al*., 2010), even minor perturbations of growing conditions attributable to positional effects within a controlled growing environment has the potential to introduce confounding variation in methylation patterning. One way of dealing with spatial variation, if it cannot be prevented, is to deploy an appropriate experimental design in order to distinguish treatment from positional effects (Brien *et al*., 2013; Cabrera-Bosquet *et al*., 2016). Experimental design normally accounts for such variability by combining blocking and randomization, along with appropriate statistical analyses (Addelman, 1970; Ruxton and Colegrave, 2011). Despite the usefulness of this approach, experimental design cannot entirely remove environmental variability (microclimate). This presents a potential challenge when attempting to link changes in DNA methylation to environmental stimuli. It is difficult to discriminate between the so-called stochastic methylation and position-dependent methylation due to the capacity of plants to promptly sense and epigenetically respond to subtle variation in ambient conditions (Gutzat and Mittelsten Scheid, 2012; Meyer, 2015).

In the present study, we combine Methylation-Sensitive Amplified Polymorphism (MSAP) and phenotypic analyses to assess the effect of microclimate on DNA methylation of barley plants growing under greenhouse conditions. Nine spring barley varieties were grown in a randomized plot design under mild soil salt stress or control conditions. Environmental, phenotypic and DNA methylation data collected at two time points are used to explore whether stochastic epigenetic may be linked to trivial environmental fluctuations. We also explore how phenotypic variability observed in these experiments correlates with differences in DNA methylation patterns.

## 2 Materials and Methods

### 2.1 Plant material and experimental design

Nine varieties of spring barley (Table 1) were grown in a controlled temperature greenhouse at the Plant Accelerator^®^ (Australian Plant Phenomics Facility (APPF), Waite Campus, University of Adelaide, Australia) from 26 June to 12 October 2013. Varieties with similar flowering times (Menz, 2010) were selected to minimize discrepancies in sampling times between varieties. The experiment comprised eight randomized blocks with two plants of the same variety per plot (Figure 1). Three seeds were sown in white pots (20 cm height × 15 cm diameter, Berry Plastics Corporation, Evansville, USA) containing 2400 g potting mixture (composed of 50% UC (University of California, Davis) potting mix, 35% coco-peat and 15% clay/loam (v/v)). Seedlings were thinned to one seedling per pot 2 weeks after sowing. Two soil salt treatments (0 mM and 75 mM NaCl (‘control’ and ‘salt stress’, respectively, hereafter) were applied to three-leaf stage seedlings (25 days after sowing (DAS)), using the protocol described by Berger *et al*. (2012). Pots were watered every 2 days for up to 60 days after sowing to 16.8% (g/g) gravimetric water content, corresponding to 0.8 × field capacity. From day 61 after sowing, plants were watered daily to 16.8% (g/g) until seed set. Leaf samples (50-100 mg) were taken for DNA extraction from blocks 1, 3, 4, 6 and 8 (Figure 1) at two time points, *viz*.: 4^th^ leaf blade after full emergence (15 days after salt treatment and 40 DAS) and flag leaf blade from the primary tiller at anthesis (62 days after salt treatment and 87 DAS). Samples were immediately snap frozen in liquid nitrogen and stored at -80 °C until DNA extraction. Whole plants were harvested at maturity and above-ground biomass was dried and weighed.

**Table 1:** List and description of barley genotypes used in this study

| No | Variety | Earliness | Year* of release | Pedigree* |  |
| --- | --- | --- | --- | --- | --- |
|  |  |  |  | Parent 1 | Parent 2 |
| 1 | Barque 73 | 6 | 1997 | Triumph | Galleon |
| 2 | Buloke | 5 | 2005 | Franklin/VB9104 | VB9104 |
| 3 | Commander | 5 | 2009 | Keel/Sloop | Galaxy |
| 4 | Fathom | 6 | 2011 | NA | NA |
| 5 | Flagship | 5 | 2006 | Chieftan/Barque | Manley/VB9104 |
| 6 | Hindmarsh | 6 | 2007 | Dash | VB9409 |
| 7 | Maritime | 6 | 2004 | Dampier/A14//Krisna/3<br>/Clipper | M11/4/DampierA14//Krisna/3<br>/Dampier/A14//Union |
| 8 | Schooner | 5 | 1983 | Proctor/PrioA (WI2128) | Proctor/CI3578 (WI2099) |
| 9 | Yarra | 5 | 2005 | VB9018/Alexis/VB9104 | NA |

Earliness to flowering score is based on a 0-9 scale, with 0 indicating very late varieties and 9 very early ones (SARDI, 2015). \*Year of release and pedigree after Menz (2010), NA = not available.

**Table 2:**
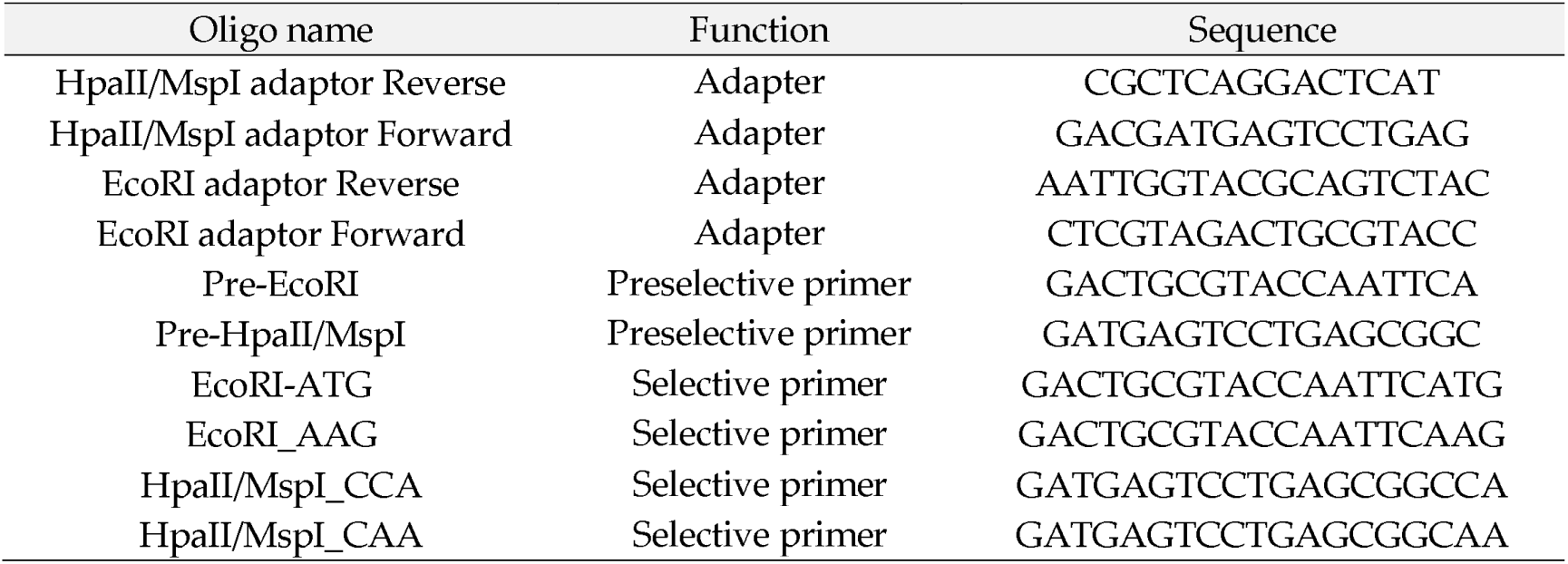
Adapter and primer sequences used for the MSAP (Rodríguez López et al., 2012).

**Figure 1:**
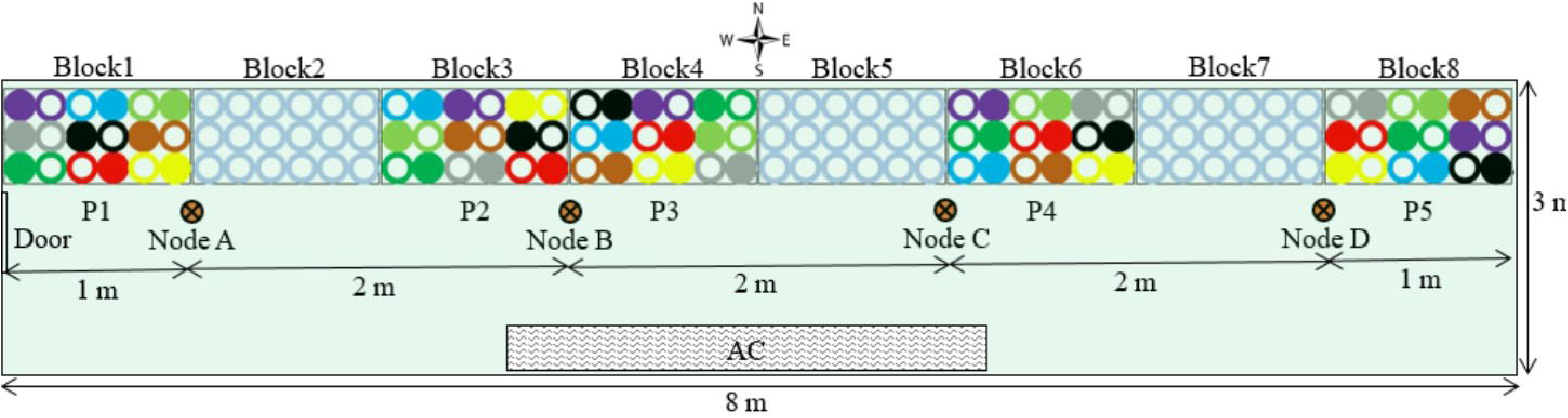
Experimental layout and plan of the greenhouse (24 m^2^). Blocks 1, 3, 4, 6 and 8 were used in this study and are respectively assigned to positions P1 to P5. Blocks 2, 5, and 7 contained empty pots. Four sensor-nodes (Node A, B, C, and D) were placed along benches, 2 metres apart and one metre from the East and West walls. Circles represent plant position in the block: hollow circles are control plants (0 mM NaCl) and full circles are treated plants (75 mM NaCl). Colours indicate barley varieties: 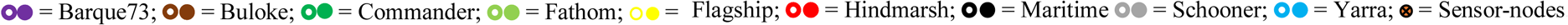. AC = air conditioning unit.

### 2.2 Greenhouse environmental conditions

The experiment was conducted in a 24 m^2^ greenhouse (∼8 m x 3 m), with a gable roof 4.5 m above the floor at the lowest and 6 m at the highest point. The greenhouse (34°58’16 S, 138°38’23 E) was oriented West-East (Figure 1). To investigate the possible causes of position dependent variability of barley response across the greenhouse, environmental factors (temperature, relative humidity and photosynthetic active rate) were recorded during the same period of the year (26 June to 12 October 2015), using four sensor-nodes located along the benches (Figure 1). Based on this period of the year, we deemed daytime to be between 7 AM and 6 PM.

The sensor-nodes were positioned 2 metres apart and 1 metre from the east and west walls (Figure 1). Each node had a combination of sensors for photosynthetic active radiance (PAR) (model Quantum, LI-COR, Lincoln, Nebraska, USA) and for humidity/temperature (Probe HMP60, Vaisala INTERCAP^®^, Helsinki, Finland). Environmental data was recorded every minute for the duration of the experiment using wireless data loggers (National Instruments, Sydney, New South Wales, Australia). Before use for further analyses, recorded data were quality controlled to remove time slots when data were not present for all four nodes. To show the overall daily fluctuation of environmental factors between sensor-nodes during the experiment, the average measure of each factor per hour was plotted for each node. Then, the vapour pressure deficit (VPD) for each time point was calculated according to Murray (1967):

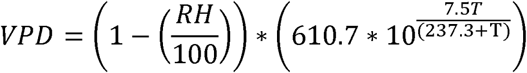

Where RH = relative humidity, T = temperature, and the factor 610.7*10^7.5T/(237.3+T)^ = saturated vapour pressure (SVP).

Pairwise comparisons of each environmental factor at sensor-node positions were performed using the Wilcoxon signed-rank test (Wilcoxon, 1945), on the R package “*ggpubr*” (Kassambara, 2019). These comparisons were performed independently for day and night periods.

### 2.3 DNA extraction

Frozen plant material was homogenized in a bead beater (2010-Geno/Grinder, SPEX SamplePrep®, USA) prior to DNA extraction using a Qiagen DNeasy kit according to the manufacturer’s instructions. DNA samples were then quantified in a NanoDrop® 1000 Spectrophotometer (V 3.8.1, ThermoFisher Scientific Inc., Australia) and concentrations were standardized to 10 ng/µl for subsequent MSAP analyses.

### 2.4 MSAP

#### 2.4.1 DNA restriction and adapter ligation

MSAP was used for the DNA methylation profiling of barley plants according to the method of Rodríguez López *et al*. (2012). To ensure marker reproducibility, DNA samples were analysed in two technical replicates. Thus, samples were digested using a methylation insensitive restriction enzyme EcoRI in combination with either *Hpa*II or *Msp*I (isoschizomers), which show differential sensitivity to cytosine methylation at CCGG positions. Digested DNA fragments were ligated to adapters (Table 1) with one end cohesive with restriction products generated by *Eco*RI or *Hpa*II/*Msp*I. Digestion and ligation reactions were performed in a single solution of 11 µl comprising: 1.1 µl T4 ligase buffer; 0.1 µl *Hpa*II; 0.05 µl *Msp*I; 0.25 µl *Eco*RI; 0.05 µl T4 ligase; 0.55 µl BSA; 1.1 µl NaCl; 1 µl Adapter *Eco*RI; 1 µl Adapter *Hpa*II/*Msp*I; 5.5 µl DNA sample and 0.3 µl pure water. Enzymes and buffer were acquired from New England Biolabs, Australia (NEB) and oligos were produced at Sigma-Aldrich, Australia. The solution was incubated for 2h at 37°C, then enzymes were inactivated at 65°C for 10 min.

#### 2.4.2 PCR

Two PCR amplifications were performed using products of the restriction/ligation reaction. First, a pre-amplification PCR was performed, in which primers complementary to adaptors but with 3’ overhangs for a unique nucleotide (*Hpa*II*/Msp*I primer +C and *EcoR*I primer +A, Table 1) were used in a pre-optimised PCR master mix (BioMix™, Bioline, Meridian Bioscience; Australia) following the manufacturer’s instructions. DNA digestion/ligation product (0.5 µl) was used for PCR amplification, with the following profile as per Rois *et al*. (2013): 72□ for 2 min, 29 cycles of 30 s denaturing at 94□, 30 s annealing at 56□ and 2 min extension at 72□, ending with 10 min at 72□ to ensure completion of the extension.

Pre-amplification products were quality assessed by 1% w/v agarose electrophoresis (80V for 2 h), before performing the selective amplification using two selective primer combinations, *Eco*RI_AAG vs. *Hpa*II/*Msp*I_CCA and *Eco*RI-ATG vs. *Hpa*II/*Msp*I_CAA. Amplified fragment detection through capillary electrophoresis was facilitated by labelling *Hpa*II/*Msp*I selective primers with the 6-FAM reporter molecule (6-CarboxyFluorescein). Just 0.3 µl of pre-amplification product was used in the pre-optimised PCR master mix and the PCR was performed as follows (Rois *et al*., 2013); 94□ for 2 min, 12 cycles of 94□ for 30 s, 65□ (and decreasing by 0.7□ each cycle) for 30 s, and 72□ for 2 min, followed by 24 cycles of 94□ for 30 s, 56□ for 30 s, and 72□ for 2 min, ending with 72□ for 10 min.

#### 2.4.3 Capillary electrophoresis

The products of the selective PCR were fractionated by capillary electrophoresis on an ABI PRISM 3730 (Applied Biosystems, Foster City, California, USA) at the Australian Genome Research Facility Ltd (Adelaide, Australia). For this, 2 µl of selective PCR products were first combined with 15 µl of HiDi formamide (Applied Biosystems) and 0.5 µl of GeneScan™ 500 ROX™ Size Standard (Applied Biosystems). The mixture was then denatured at 95□ for 5 min and snap-cooled on ice for 5 min before sample fractionation at 15 kV for 6 s and at 15 kV for 33 min at 66□.

#### 2.4.4 MSAP data analysis

MSAP profiles obtained using *Hpa*II and *Msp*I were used to generate; 1) a qualitative binary matrix of allelic presence/absence scores, and 2) a quantitative matrix of allelic peak height using GeneMapper Software v4 (Applied Biosystems). Qualitative epigenetic changes associated with greenhouse positional effect were analysed using fragment sizes between 100 and 550 base pairs, which were selected to estimate epigenetic distance between individual plants (EpiGD) and subpopulations of plants (PhiPT) and perform Principal Coordinate Analyses (PCoA), using GenAlex 6.501 (Peakall and Smouse, 2012).

Quantitative analysis of peak height was used to examine the effect of position on the methylation status of individual loci. We searched for MSAP markers that were differentially methylated between experimental blocks by comparing the fragment peak heights to survey for position effects on the plant methylation profile (Rodríguez López *et al*., 2012). Before differential methylation analysis, model-based normalization factors were calculated for the peak height libraries using the weighted trimmed mean method of Robinson and Oshlack (2010). For each variety and sampling method, peak heights were extracted and analysed individually using the modelling approach of McCarthy *et al*. (2012). To ensure the peak heights could be compared between positions, the individual models contained a term to account for variation between blocks as well as a term to capture the differences between the control and salt stress treatments. A likelihood ratio test was then performed to determine whether estimated coefficients for the positions were equal (McCarthy *et al*., 2012). The p-values from these tests were then adjusted for multiple comparisons using the false discovery rate method of Benjamini and Hochberg (1995). Analyses were conducted using the R package *edgeR* (Robinson *et al*., 2010), in the R statistical computing environment (R Core Team, 2019).

The extent of epigenetic divergence between salt treatments at the two developmental stages (4^th^ leaf and anthesis) was assessed, first by performing a multiple correspondance analysis (MCA) on MSAP marker data. A linear discriminant analysis (LDA) was then performed on the MCA results. These analyses, refered to as MC-LDA thereafter, were done using the R packages FactoMineR and MASS (Lê *et al*., 2008; R Core Team, 2019). To visualise the results of comparisons involving more than two groups, the first two linear discriminant factors (LD1 and LD2) were plotted. Otherwise, a density plot of LD1 was performed.

### 2.5 Assessment of correlations between epigenetic profiles and plant phenotype

Epigenetic and phenotypic variability were estimated using averaged data per position for all nine barley varieties (Bishop *et al*., 2015). The software GraphPad Prism 6 v008 (Graph-Pad Software, San Diego, California, USA) was used to perform statistical analyses. Values of above-ground plant biomass were normalized by computing the ratio of plant biomass over the mean biomass for each individual experiencing the same treatment across all positions. The same formula was applied to grain yield. This normalization was intended to address quantitative variability between treatments and among barley genotypes. Then, biomass and yield distance matrices were generated using the difference between normalized values of any two individual plants.

We performed a Mantel Test (Mantel, 1967) to estimate the significance of the correlations between epigenetic distance and plant biomass, and position in the greenhouse. For this, we used matrices generated from epigenetic distance, physical distance and phenotypic (biomass or yield) differences estimated as described above. In all cases, the level of significance of the observed correlations was tested using 9,999 random permutations. Since both enzymes (*Hpa*II, *Msp*I) are methylation sensitive (Walder *et al*., 1983; Reyna-López *et al*., 1997), these enzymes can independently show epigenetic marks across the genome. Therefore, our inferences about plant epigenetic profile thereafter relate to results obtained using either enzyme or a combination of both.

## 3 Results

### 3.1 Microclimatic variability in the greenhouse

Data quality control of climatic data provided 47,144 and 54,983 time-points of data recording for the periods of day and night, respectively. These correspond to time-points when recording was obtained simultaneously in all sensor-nodes. There was clear evidence of both spatial and temporal variation for temperature, photosynthetically active radiation (PAR) and relative humidity (RH) within the experimental area (Figures 2 and 3).

**Figure 2:**
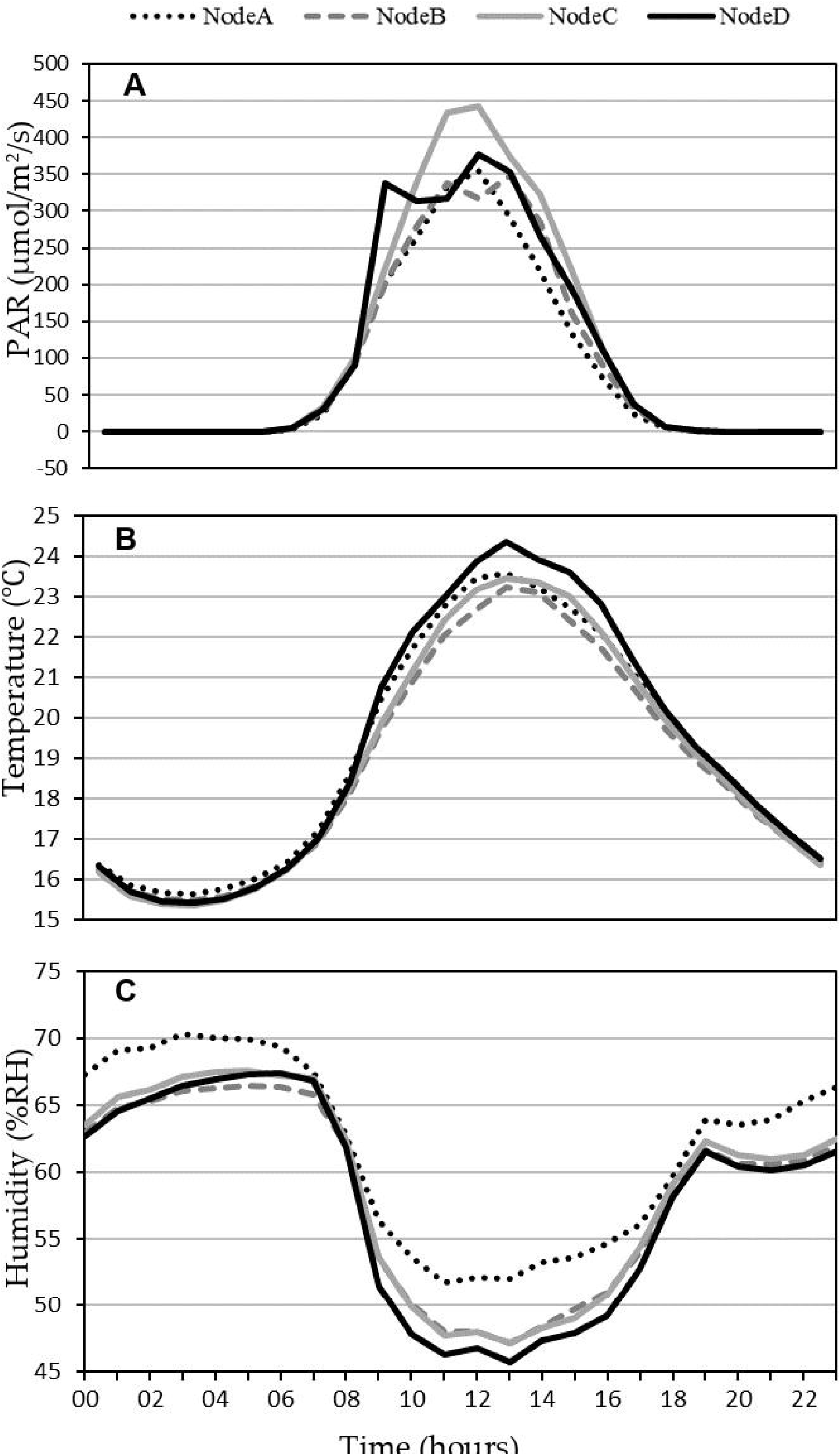
Average daily fluctuations of climatic conditions in the greenhouse. (A) light, (B) temperature and (C) relative humidity were recorded over the period from 26 June to 12 October 2015, at four positions (Node A-D from West to East) in the greenhouse.

**Figure 3:**
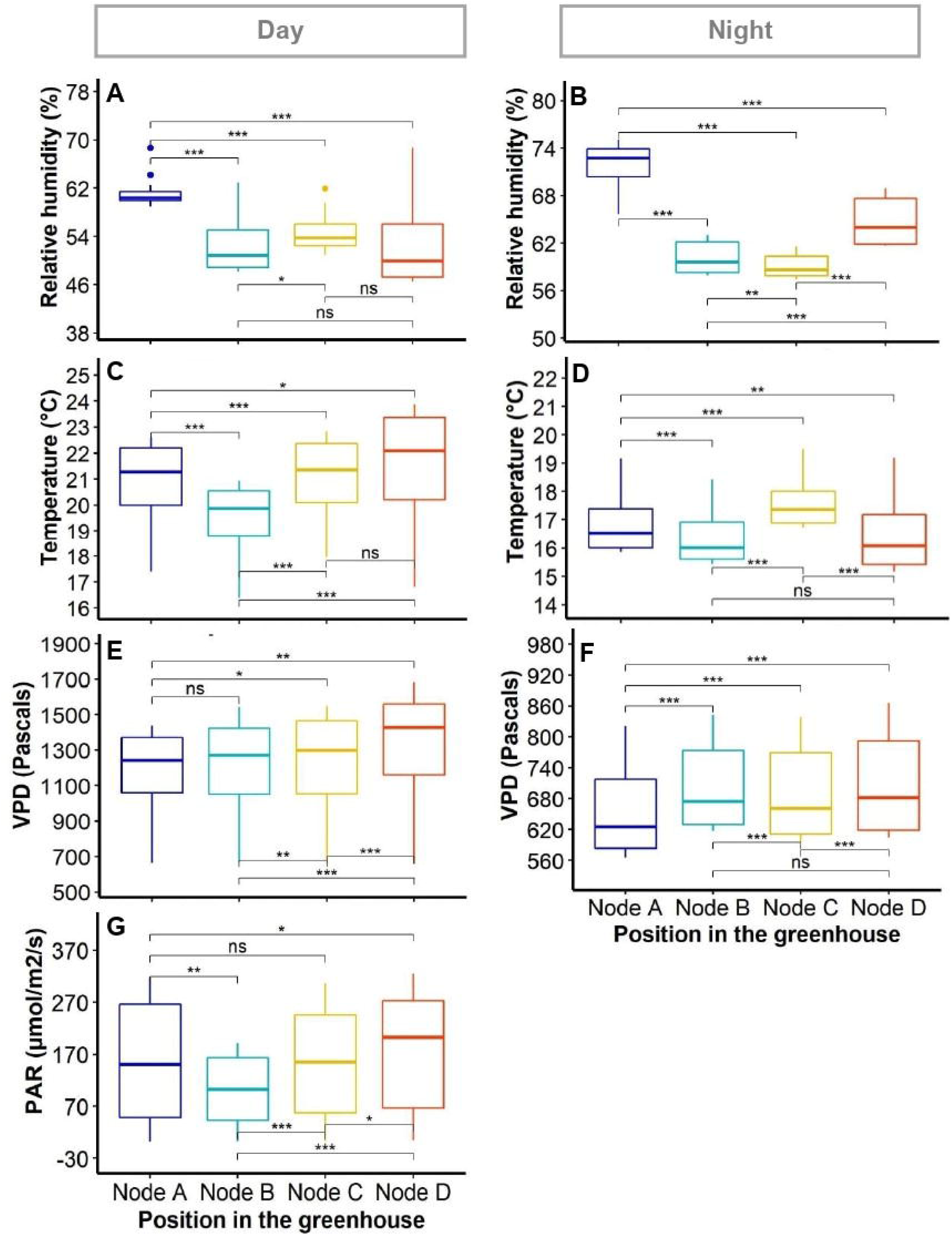
Variability of climatic factors in the greenhouse. The boxplots show variations within positions and compare data between sensor-nodes based on Wilcoxon paired signed-test. Asterisks (*), (**) and (***) indicate the significance of the difference between positions (nodes) for P-value < 0.05, 0.01 and 0.001, respectively; ns = difference not significant. The PAR was deemed as null at night.

The average dynamics of climatic data in the greenhouse showed a higher PAR between 8 AM and 10 AM at the East side than the rest of the greenhouse (node D, Figure 1). The PAR was also variable during the day between node positions, with sensor-node B (Centre-West, Figure 1) recording the lowest PAR values around 12 PM (Figure 2A). The average temperatures evolved broadly in the same way at all node positions, with only around 1.5□ difference between the most divergent nodes at the warmest time of day (Figure 2B). The RH was the highest at node A (West side of the greenhouse, Figure 1) during both day and night, and was significantly different from the rest of the positions during the day (Figures 2C and 3). The node D (East end of the greenhouse) presented the lowest RH during the day; it was not significantly different from nodes B and C (Figure 3A).

Although there was no clear evidence of gradient between sensor-nodes for any of the climatic factors (i.e. RH, temperature, VPD and PAR, the pairwise comparison of data from sensor-nodes using Wilcoxon paired signed-rank test showed significant differences between positions for each variable (Figure 3A-G). Such differences were present during both day and night periods in the greenhouse. The RH appeared particularly variable at night between all positions of sensor-nodes (Figure 3B).

### 3.2 Correlation between DNA methylation profile and plant position in the greenhouse

Plant DNA methylation profiles derived from MSAP data generated 269 alleles with sizes between 100 and 550 base pairs across samples from all nine barley varieties. PCoA of MSAP profiles for barley variety at anthesis showed grouping of samples more by plant position than salt treatment, regardless of the enzyme combination used (Figures 4A and B). The first coordinate Eigen space matched with the position of the plants in the greenhouse in the West-East direction (Figure 4). The Mantel test using all treatment samples together showed weak correlations between plant epigenetic profiles and plant positions in the greenhouse at 4^th^ leaf stage, and more significant corrections at anthesis (Table 3). For instance, for the variety Schooner, the Mantel test between pairwise epigenetic distance and plant position at the 4^th^ leaf stage of barley development resulted in weak correlations for both *Hpa*II (R^2^ = 0.11, P-value = 0.025, Figure 5A) and *Msp*I (R^2^ = 0.12, P-value < 0.022, Figure 5C). Apart from two varieties (Buloke and Schooner), none of the remaining varieties showed a significant correlation between position and epigenetic profile at the 4^th^ leaf stage (Table 3, Figures S1). Conversely, these correlations were stronger at anthesis for the same variety Schooner (R^2^ = 0.48 and R^2^ = 0.45, for *Hpa*II and *Msp*I, respectively, Figure 5B and D), with greater significance of the P-values (0.001). Additionally, all the remaining varieties showed significant correlation (P-value at least < 0.05) between DNA methylation profile at anthesis and the plant position in the greenhouse (Table 3; Figure S1). The correlations at anthesis were high (R^2^ > 0.3) for all varieties, except Buloke and Maritime (Table 3).

**Table 3:** Correlation between pairwise epigenetic distance and physical distance. Nine barley varieties were used, comprising ten individuals per variety, five replicates for control and stress plants. Samples were collected from the 4^th^ leaf (at 4^th^ leaf stage) and flag leaf (at anthesis). Epigenetic distances correspond to the Phi statistics of the MSAP markers between plant individuals. The coefficient of determination (R^2^) was calculated according to Mantel (1967) using GenAlex 6.5. Asterisks (*), (**) and (***) indicate significant correlation between treatments for P-value < 0.05, 0.01 and 0.001, respectively, estimated based on 9999 permutations.

| Varieties | Coefficient of determination ($R^2$) | | | |
| --- | --- | --- | --- | --- |
|  | *HpaII* |  | *MspI* |  |
|  | 4th leaf | Anthesis | 4th leaf | Anthesis |
| Barque73 | 0.003 | 0.320** | 0.010 | 0.315 |
| Buloke | 0.103* | 0.001 | 0.059 | 0.220* |
| Commander | 0.052 | 0.332** | 0.050 | 0.332** |
| Fathom | 0.038 | 0.425**** | 0.079* | 0.527**** |
| Flagship | 0.038 | 0.451*** | 0.001 | 0.214* |
| Hindmarsh | 0.008 | 0.305** | 0.004 | 0.233* |
| Maritime | 0.014 | 0.130* | 0.071* | 0.144* |
| Schooner | 0.112* | 0.476*** | 0.120* | 0.447*** |
| Yarra | 0.002 | 0.147* | 0.027 | 0.385* |
| Average | 0.041 | 0.287 | 0.047 | 0.313 |

**Figure 4:**
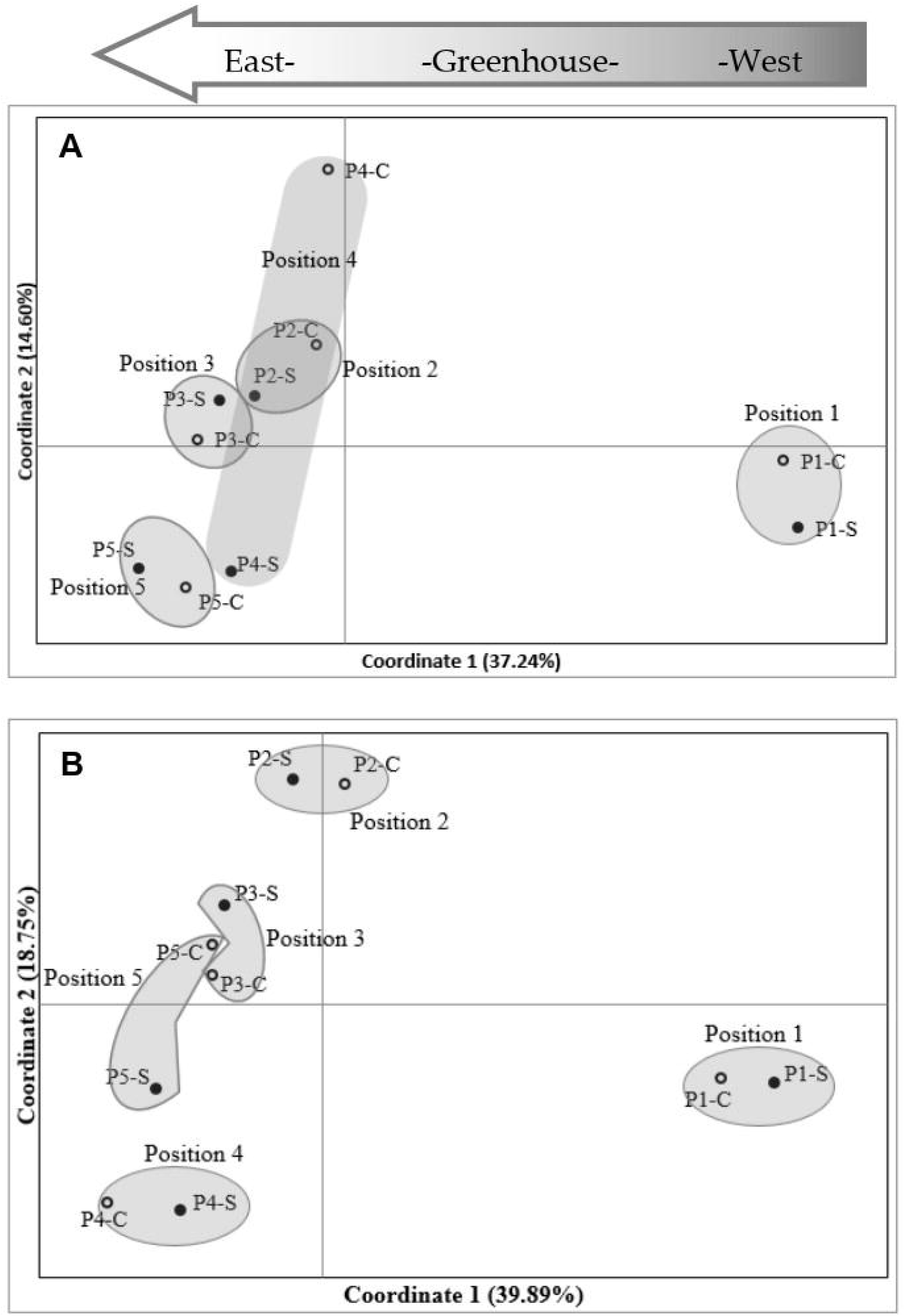
Principal coordinates analysis (PCoA) of MSAP (methylation sensitive amplified polymorphism) markers in barley variety Commander. MSAP markers were generated using five replicates of control (0 mM NaCl) and stress (75 mM NaCl) plant samples, for *Hpa*II (A) and *Msp*I (B). Positions 1 to 5 indicate experimental block numbers; Symbols filled in black and hollow symbols represent salt stress (-S) and control (-C) samples, respectively. The PCoAs show sample distribution in the first two principal coordinates. Numbers in brackets represent the proportion of variation explained by the coordinate.

**Figure 5:**
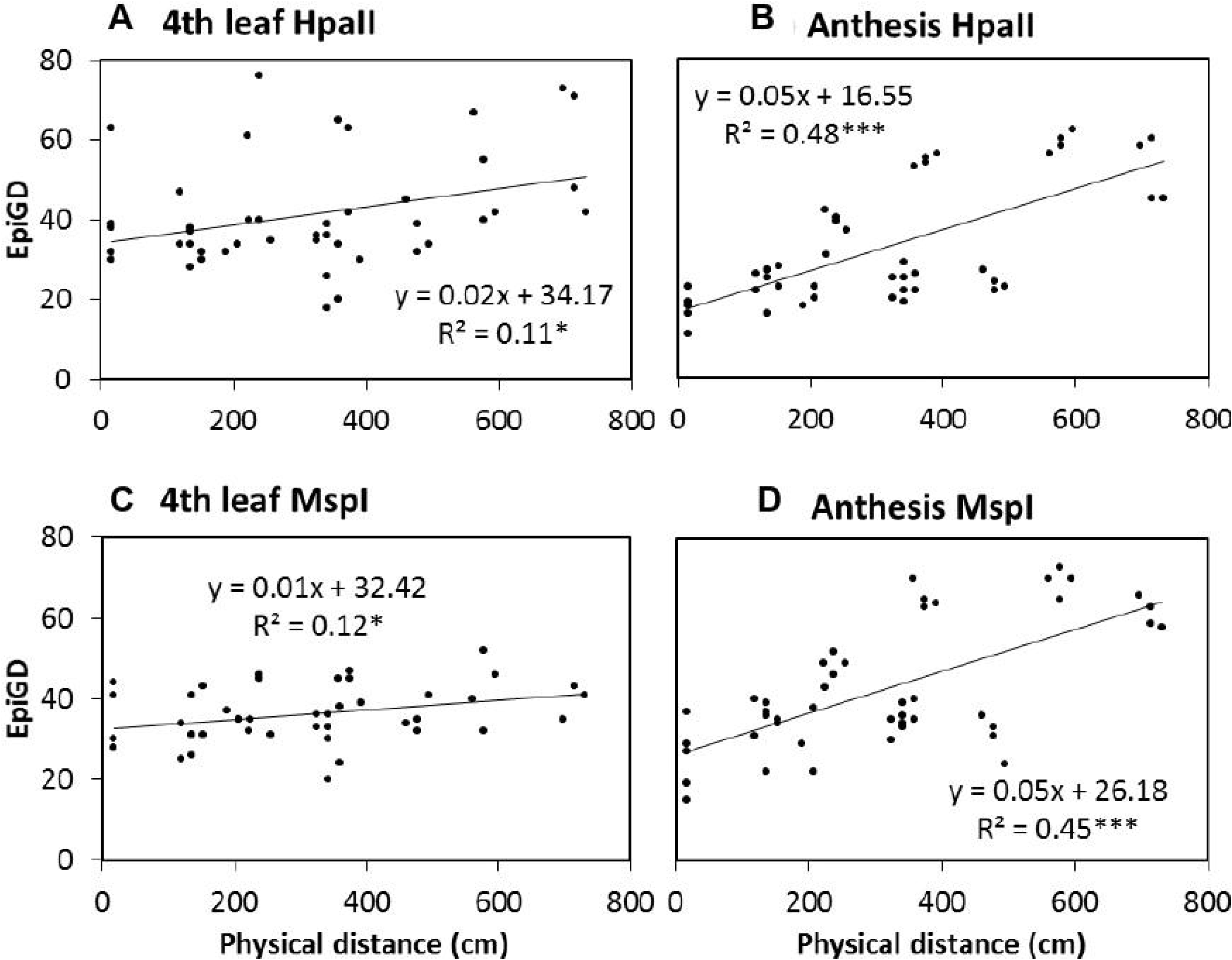
Correlation between pairwise epigenetic distance (Epi GD) and plant position in the greenhouse. The epigenetic distance was estimated at 4^th^ leaf stage (a, c; 40 days after sowing) and anthesis (c, d; 87 days after sowing) of barley variety Schooner, using *Hpa*II (a, b) and *Msp*I (c, d) for the MSAP (methylation sensitive amplified polymorphism) analysis. Five replicates of control (0 mM NaCl) and stress (75 mM NaCl) were analysed together and dots represent pairwise comparisons between individual plants. Equations represent the formula of the regression line, R^2^ represents the coefficient of determination, calculated according to Mantel (1967) using GenAlex 6.5. Asterisks (*) and (***) indicate significant correlation between treatments for P-value < 0.05 and 0.001, respectively, estimated based on 9999 permutations.

The comparison of peak heights of MSAP markers generated from plants growing in different positions revealed significant differences between positions for some alleles (Figure 6). In general, significant differences in peak height were observed between plants in position P1 and the other positions (Figure 6). Overall, peak heights showed logarithmic trends (both positive and negative), significantly associated with the West-East distribution of the samples. A few markers were significantly different in peak heights over all positions (Table 4).

**Table 4:** List of salt-induced methylation marker alleles showing significant peak height differences between the five experimental blocks. logFC = log fold change; logCPM = log counts per million; LR = likelihood ratio statistics; FDR = false discovery rate.

| Variety | Sample tissue | Enzyme/Primer | allele | logFC | logCPM | LR | PValue | FDR |
| --- | --- | --- | --- | --- | --- | --- | --- | --- |
| Barque73 | Flag leaf | *HpaII*/ATG-CAA | 403.76 | 0.884 | 12.895 | 12.082 | 0.001 | 0.019 |
| Barque73 | Flag leaf | *HpaII*/ATG-CAA | 221.61 | -1.749 | 14.043 | 9.817 | 0.002 | 0.032 |
| Flagship | 4th leaf | *HpaII*/ATG-CAA | 221.61 | -1.202 | 13.901 | 10.507 | 0.001 | 0.036 |
| Yarra | Leaf before flag | *HpaII*/ATG-CAA | 361.55 | -0.653 | 12.238 | 10.505 | 0.001 | 0.036 |
| Yarra | Leaf before flag | *HpaII*/ATG-CAA | 167.6 | -0.796 | 12.866 | 8.726 | 0.003 | 0.040 |
| Yarra | Leaf before flag | *HpaII*/ATG-CAA | 543.70 | 0.816 | 12.508 | 8.286 | 0.004 | 0.040 |

**Figure 6:**
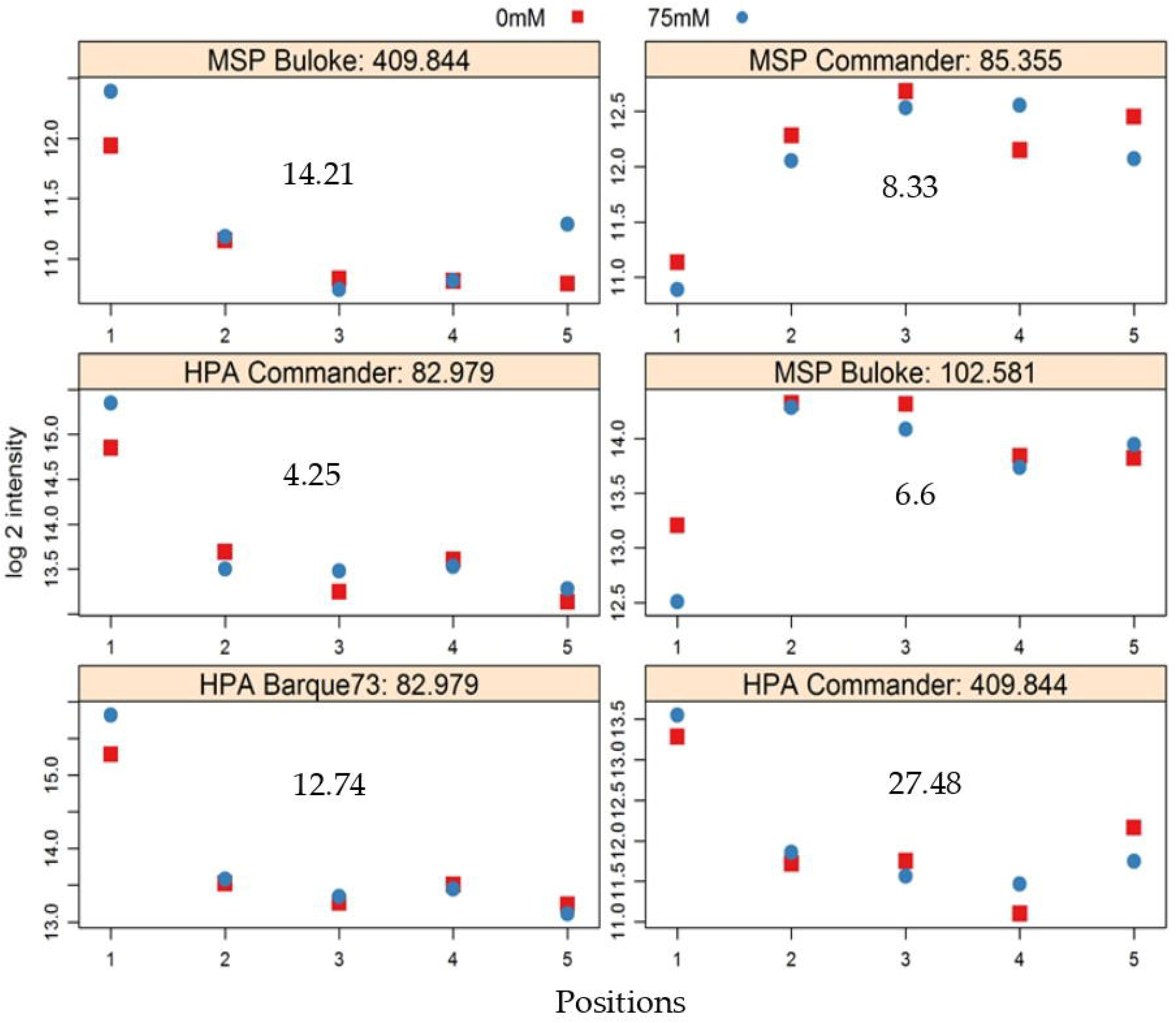
Exemplars of MSAP (methylation sensitive amplified polymorphism) alleles that show significant differences in peak height between positions in the greenhouse. Markers were detected in control (0 mM NaCl, red symbols) and stress (75 mM NaCl, blue symbols) plants; Vertical axis shows logarithm 2 (log 2) of peak height intensity and the horizontal axis represents positions in the greenhouse, in the West to East direction. The grey number in each plot represents -log10 of p-values. The title of each plot shows the enzyme used (either *Hpa*II (HPA) or *Msp*I (MSP), the variety, and the allele identity number.

However, positional effect did not thwart the ability to differentiate between salt-stressed and control plants. The MC-LDA on MSAP marker data was able to separate salt stressed plants from those given control conditions (Figures 7A-B). Furthermore, epigenetic divergence between treatment groups increased with time, with control and stress plants consistently more similar at the 4^th^ leaf stage than at anthesis across all varieties (Figures 7 A-B and S2). MC-LDA of salt treatments could nevertheless discriminate treatments at both stages even though epigenetic divergence was strongly influenced by developmental stage (Figures 7 C and S2).

**Figure 7:**
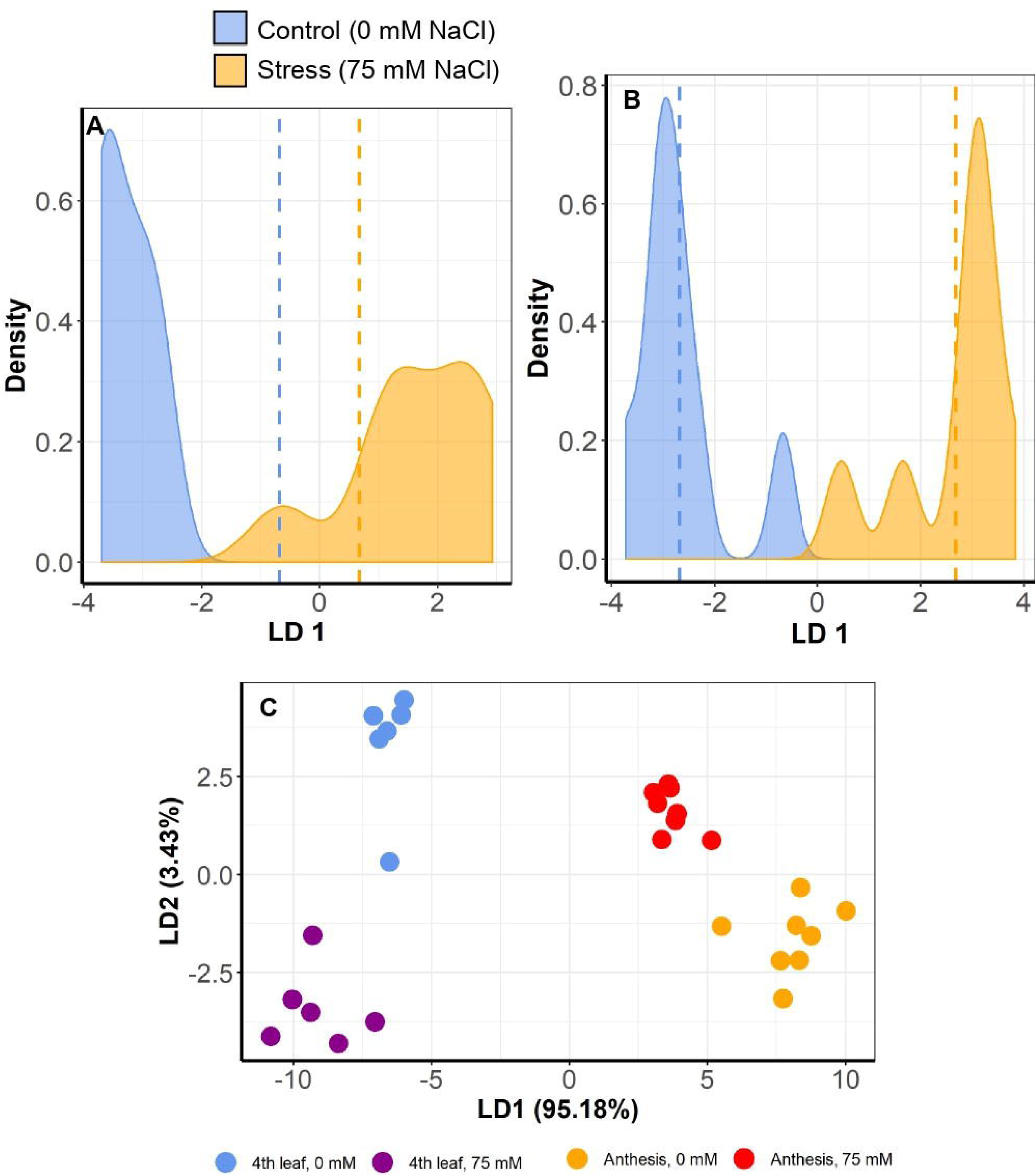
Multiple Correspondence and Linear Discriminant Analyses (MC-LDA) of MSAP markers in barley variety Commander under salt stress (75 mM) and control (0 mM) conditions. The panel shows density plots of LD function between stress and control plants, at 4th leaf stage (A) and at anthesis (B). Dashed vertical lines represent the mean LD1 in 2 groups’ comparisons. The graph C shows MC-LDA plots comparing the salt treatment groups at both 4th leaf and anthesis stages. Similar plots for the other varieties are presented in supplementary Figure S2.

### 3.3 Correlations between barley phenotype, epigenome and position

There was a clear trend in the final biomass of all nine barley varieties according to position, with a progressive increase from position P1 (West side of the greenhouse) to position P5 (East side) (Figure 8A). This relationship was a logarithmic trend, both in the control and stressed plants. The average grain yield of the barley varieties showed the same West-East trend as the biomass (Figure 8B). However, when varieties were examined separately, both logarithmic and polynomial trends were observed (Figure S3).

**Figure 8:**
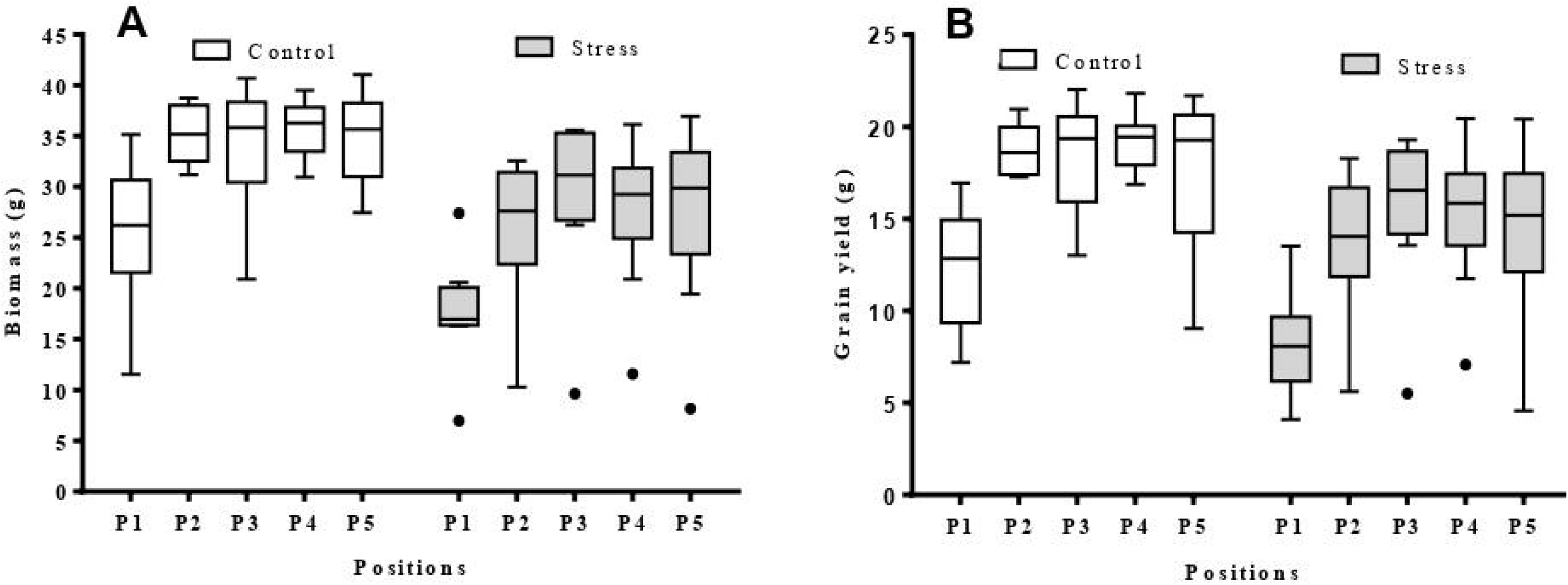
Box plots showing biomass and grain yield range per position (P1-5) in the greenhouse (n = 9). (a) biomass per position for control and stress plants; (b) grain yield per position for control and stress plants; The average data was obtained from nine barley varieties (Barque 73, Buloke, Commander, Fathom, Flagship, Hindmarsh, Maritime, Schooner and Yarra).

Assessment of the relationship between pairwise differences in epigenetic distance and in grain yield showed significant correlations (P-values < 0.05) in control plants of six of nine varieties (Buloke, Commander, Fathom, Maritime, Schooner, Yarra), with R^2^ varying between 0.247 and 0.907 (Table 5; Figure S4). Likewise, stress plants showed significant correlations (P-values at least < 0.05) between grain yield and methylation profile in six varieties (Barque 73, Buloke, Commander, Flagship, Maritime, Schooner), with R^2^ between 0.164 and 0.921 (Table 5; Figure S4). An example of significant correlations between grain yield and epigenetic distance is presented in Figure 9A-D, for the variety Schooner.

**Table 5:**
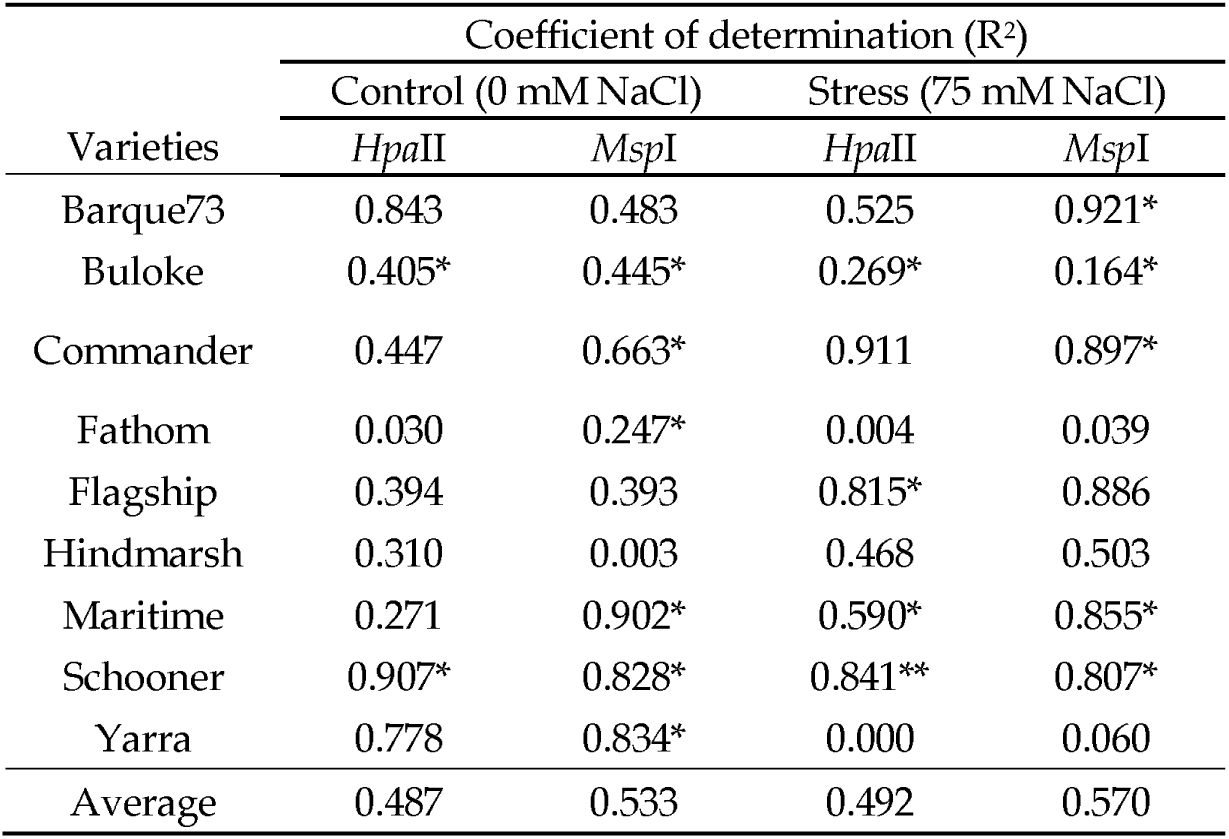
Correlation between epigenetic distance and grain yield of nine barley varieties. Epigenetic distance between plants was calculated based on MSAP data generated using *Hpa*II and *Msp*I. Coefficients of determination (R^2^) were computed according to Mantel (1967) using five replicates for each treatment per variety. Asterisks (*) and (**) indicate significant correlation between treatments for P-value < 0.05, and 0.01, respectively, estimated based on 9999 permutations.

| Varieties | Coefficient of determination ($R^2$) | | | |
| --- | --- | --- | --- | --- |
|  | Control (0 mM NaCl) |  | Stress (75 mM NaCl) |  |
|  | *HpaII* | *MspI* | *HpaII* | *MspI* |
| Barque73 | 0.843 | 0.483 | 0.525 | 0.921* |
| Buloke | 0.405* | 0.445* | 0.269* | 0.164* |
| Commander | 0.447 | 0.663* | 0.911 | 0.897* |
| Fathom | 0.030 | 0.247* | 0.004 | 0.039 |
| Flagship | 0.394 | 0.393 | 0.815* | 0.886 |
| Hindmarsh | 0.310 | 0.003 | 0.468 | 0.503 |
| Maritime | 0.271 | 0.902* | 0.590* | 0.855* |
| Schooner | 0.907* | 0.828* | 0.841** | 0.807* |
| Yarra | 0.778 | 0.834* | 0.000 | 0.060 |
| Average | 0.487 | 0.533 | 0.492 | 0.570 |

**Figure 9:**
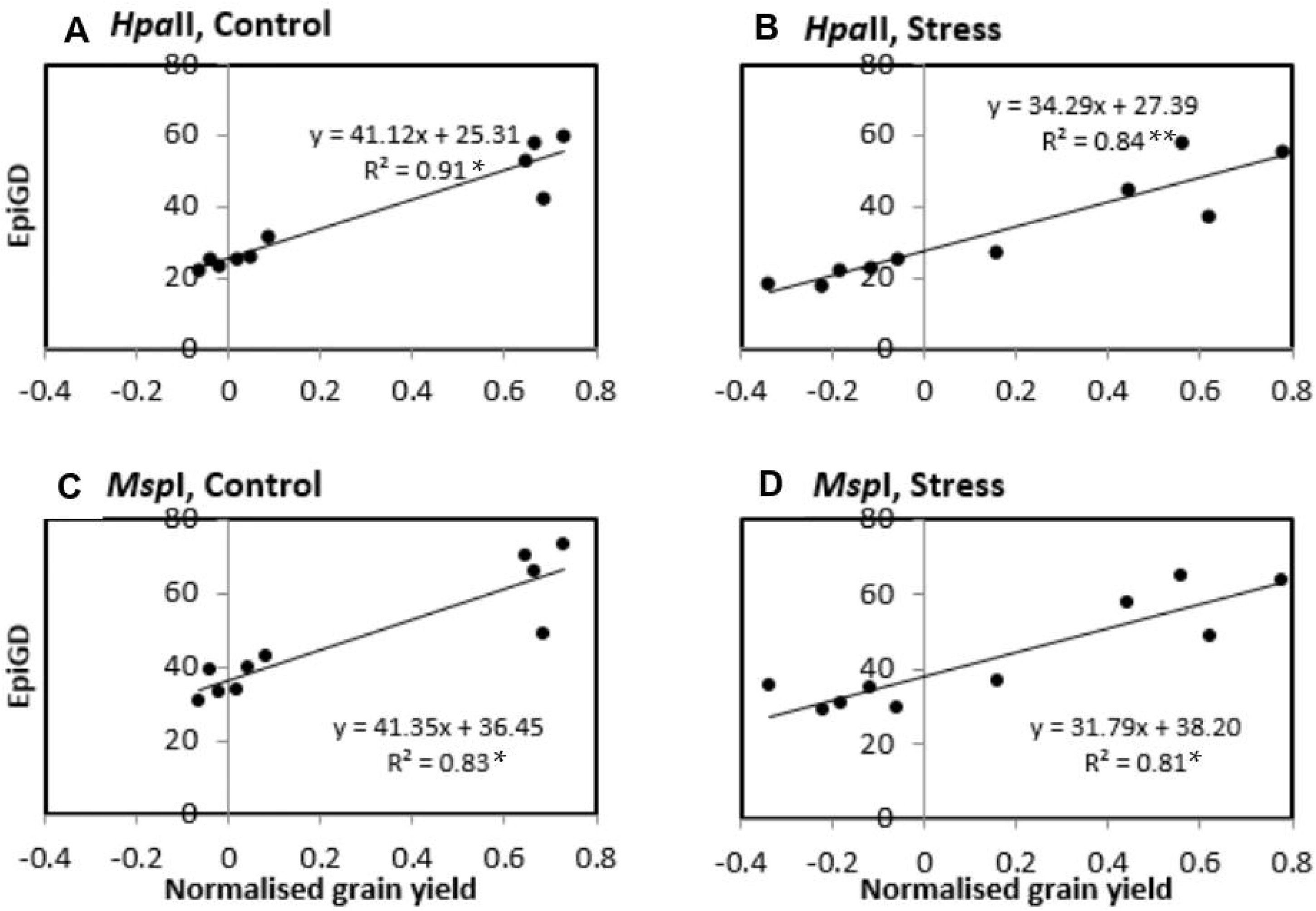
Correlation between pairwise epigenetic distance (EpiGD) and pairwise difference in grain yield between plants of the variety Schooner. The correlation was tested according to Mantel (1967) using GenAlex 6.5. Epigenetic distance between plants was calculated based on MSAP (methylation sensitive amplified polymorphism) data generated using *Hpa*II (a, b) and *Msp*I (c, d). Pairwise differences in grain yield between plants were calculated separately for control (a, c) and stress (b, d) plants. Values of grain yield were normalized by computing the ratio of each individual plant grain yield over the mean grain yield for the same treatment across all positions. The dots represent pairwise comparisons between individual plants; equations represent the formulae of the regression line; R^2^ represents the coefficient of determination of the Mantel test; asterisk (*) and (**) indicate significant correlation between treatments for P-value < 0.05, and 0.01, respectively, estimated based on 9999 permutations.

## 4 Discussion

### 4.1 Stochastic DNA methylation is explained by microclimatic differences

The randomized block design aims to minimise unexplained variation between treatments, and has emerged as a preferred method in plant field trials and in controlled environment experiments (Edmondson, 1989; Guertal and Elkins, 1996; Brien *et al*., 2013). However, while block homogeneity is difficult to achieve, variability between blocks in the same experimental setting is often either ignored, attributed to randomness (Raj and van Oudenaarden, 2008; Karan *et al*., 2012; Tricker *et al*., 2012) or in the context of epigenetic research, explained by spontaneous occurrence of the methylation (Becker *et al*., 2011; Baulcombe and Dean, 2014; van der Graaf *et al*., 2015).

In this study, we found evidence suggesting that microclimatic variation within a greenhouse was sufficient to trigger variability in the plant DNA methylation profile in a manner that was both independent of the experimental treatment and greater in magnitude. The clarity of the climatic variables measured across the experimental blocks, and the associated cline in methylation patterning is suggestive that each plant experienced a unique combination of climatic factors during the experimental period, and that this induces, at least partly, changes in methylation patterning. Similar observations were also reported for other greenhouse studies (Brien *et al*., 2013; Both *et al*., 2015; Cabrera-Bosquet *et al*., 2016). This finding is inconsistent with spontaneous DNA methylation being entirely responsible for the plant-plant variability in such experiments (Becker *et al*., 2011; van der Graaf *et al*., 2015), and throws into question how best to discriminate epigenetic responses to micro-environment fluctuations from those attributable to stochastic noise. Moreover, the effect of position can easily be overlooked in snap-shot exposure experiments, since the timeframe from stress exposure to induction of position-dependent methylation markers is critical but also likely to vary between loci. Support for this reasoning can be taken from our findings that it was possible to separate salt and control samples by discriminate analysis at the 4^th^ leaf stage and at anthesis but with higher divergence at the later stage. At the same time, correlation between epigenetic differences and physical distance among plants at anthesis (87 DAS) was stronger than at the 4^th^ leaf stage (40 DAS), indicating that exposure to the stressor and positional microclimates both have a cumulative effect on the plant epigenome. These observations are congruent with the concept that plant adaptive improvisation, through DNA methylation, is proportional to the severity and duration of the environmental cue to which the plant was exposed (Soen *et al*., 2015). In this sense, the scale of the effect induced by intervention stress (salt) needs to be weighed against those imposed by coincidental stresses (microenvironment effects) but also by those associated with development or ageing, as was reported in humans (Gentilini *et al*., 2015). Any truly stochastic DNA methylation would represent residual variation. Previous studies have observed the influence of mesoclimatic conditions (Herrera and Bazaga, 2010) and factors such as temperature (Hashida *et al*., 2006), humidity (Tricker *et al*., 2012) or light (Barneche *et al*., 2014; Meyer, 2015) on methylome variability. However, the current study suggests, for the first time, that even slight variations in climatic factors (temperature, humidity or light) are sufficient to induce modifications in the plant DNA methylation profile, and that this can be sufficient to mask effects of mild stresses, as was observed here for salt stress. We certainly do not contend that all nascent methylation arises in response to environmental or biotic effectors but we do argue that far more care is needed before discounting unaccounted epigenetic variation as stochastic noise.

### 4.2 Positional effect affects salt stress-induced DNA methylation changes in barley

Positional effects in greenhouse experiments are well established and if not properly accounted for can generate uncharacterised background noise that can mask the effect of the experimental treatment (Edmondson, 1989; Guertal and Elkins, 1996; Brien *et al*., 2013). Spatial variability in coincident environmental factors has potential to introduce variability between replicate plants’ development and response to experimental treatments (Edmondson, 1989; Guertal and Elkins, 1996). Such spatial variability is liable to introduce flaws in measurements and observations between replicates that, in fact, were not experiencing exactly the same constraints (Addelman, 1970). This can compromise the search for relationships between experimentally controlled stressors (in our study, soil salt stress) and perturbations in epigenetic profiles. Indeed, in the present work the observed negative correlation between RH and differences in epigenetic differentiation between control and salt stressed pairs of plants growing in the different positions suggests that variations in environmental factors has interfered with reaction of the plant to mild salt stress. One possible mechanistic explanation is that the observed West to East decrease in RH changed the plant’s requirement for water (Barnabás *et al*., 2008; Verslues and Juenger, 2011), and this in turn may have affected the level of salt stress experienced by each plant. In this way, plants were grown under the same salt treatment but because they experienced different RH, are likely to exhibit a different response to the salt stress; hence the inconsistent salt-induced DNA methylation profiles.

### 4.3 Phenotypic differences associated to greenhouse microclimates correlate with epigenetic differences

The finding here of a plastic response by barley plants in terms of biomass and grain yield to subtle differences associated with greenhouse position corroborates earlier work by Lacaze *et al*. (2008) who suggested that barley is responsive to fluctuations in ambient conditions. We postulate that the irregularity of phenotypic variability patterns across barley varieties and treatments may have emerged from two complementary factors; 1) the genetic variability among barley varieties leading to differential responsiveness to positional effect, as reported elsewhere (Lacaze *et al*., 2008; Kren *et al*., 2015), and 2) the randomness of spatial microclimatic conditions, which did not have a linear spatial gradient. The influence of a genotype-by-environment effect on plant phenotype was expected (Gianoli and Palacio-López, 2009; Aspinwall *et al*., 2015), but the scale of phenotypic variation induced by small-scale environmental variation was not. Our findings highlight the possibility for plants to show substantial phenotypic responses to even slight variations in ambient conditions, and that homogeneity in temperature control does not have over-riding importance. Furthermore, our discovery of a significant correlation between barley MSAP profiles and grain yield suggests that DNA methylation could at least reflect and possibly contribute towards the plastic variation in plant phenotypes. These results are in accordance with a mounting body of evidence that plant plasticity is at least partly epigenetically governed (Boyko and Kovalchuk, 2008; Rois *et al*., 2013; Baulcombe and Dean, 2014; Aspinwall *et al*., 2015). Considered together, our results demonstrate a tight interplay between plant epigenome, environment and phenotype.

## 5 Conclusions

Homogeneity of environmental conditions is practically difficult to obtain in a greenhouse (Edmondson, 1989; Guertal and Elkins, 1996; Brien *et al*., 2013). Awareness of plant sensitivity to microclimate is therefore important, especially in epigenetic studies, where plant epigenomes seem to be extremely responsive to small fluctuations in environmental factors. This study reveals that at least some of the DNA methylation previously considered stochastic is likely to have been, at least partially, induced by 1) positional effects on growth conditions, 2) differences in the length of plant exposure to relatively trivial variations in environment and 3) synergistic effects of stress treatment (mild salt stress in this case) and microclimatic conditions. The correlation between phenotypic DNA methylation differentiations between plants grown in different microclimates suggests that position-induced DNA methylation, previously ignored or considered as stochastic, may be a substantial source of phenotypic variability. Accordingly, we advocate that future epigenetic analyses should take into account the effect of micro-variations in environmental factors by careful experimental design and by considering position-induced DNA methylation markers as strong candidates for finely-tuned response to small environmental changes. We also propose that further research is needed to untangle microclimate-induced epigenetic variations from epigenome instability due to experimental treatment and developmental stage.

## 6 Conflict of Interest

The authors declare that the research was conducted in the absence of any commercial or financial relationships that could be construed as a potential conflict of interest.

## 7 Author Contributions

M.K. performed the experiments, analysed the data and wrote the manuscript; J.T. performed the statistical analysis of MSAP peak heights; M.J.W., E.S.S., B.B. and C.M.R.L. conceived the experiments and supervised the work. All authors read and commented on the manuscript.

## 8 Funding

M.K. was supported by Australian Awards, AusAID (Australian Agency for International Development); M.J.W. was partly supported by the Biotechnology and Biological Sciences Research Council (BBS/E/0012843C) and C.M.R.L. is currently partially supported by the National Institute of Food and Agriculture, U.S. Department of Agriculture, Hatch Program number 2352987000.

### 9 Acknowledgments

We are grateful to AusAID (Australian Agency for International Development) for providing an Australian Awards Scholarship to MK for his PhD. The Biotechnology and Biological Sciences Research Council (BBSRC) strategic program grant (BB CSP1730/1) paid for MW time. We also acknowledge Olena Kravchuk for contributing to the experimental design, Kate Dowling for the quality control of environmental data in the greenhouse, and technical staff at The Plant Accelerator, Australian Plant Phenomics Facility, which is funded under the National Collaborative Research Infrastructure Strategy of the Australian Commonwealth.

## 10 Supplementary Material

The Supplementary Material for this article can be found online at:

**Figure S1:** Correlation between epigenetic distance (Epi-GD) and geometric distance between plants (cm, centimetre) using the Mantel test, which was performed on data from nine barley varieties (Barque 73, Buloke, Commander, Fathom, Flagship, Hindmarsh, Maritime, Schooner and Yarra) and methylation sensitive enzymes *Hpa*II (a-f) and *Msp*I (g-l). Analyses involved control and stress plants together (a, b, g and h), control plants only (c, d, i and j) or stress plants only (e, f, k and l). Correlations were generally lower at 4^th^ leaf stage (a, c, e, g, i and k) than at anthesis (b, d, f, h, j and l), indicating that positional effect is cumulative during plant development.

**Figure S2:** Multiple Correspondence and Linear Discriminant Analyses (MC-LDA) of MSAP markers in barley varieties (Barque73, Buloke, Fathom, Flagship, Hindmarsh, Maritime, Schooner and Yarra) under salt stress (75 mM) and control (0 mM) conditions. The panel shows density plots of LD function between stress and control plants, at 4th leaf stage (A, D, G, J, M, P, S, V) and at anthesis (B, E, H, K, N, Q, T, W). Dashed vertical lines represent the mean LD1 in comparisons of two groups. Graphs of panel C, F, I, L, O, R, U and X are MC-LDA plots comparing the salt treatment groups at both 4th leaf and anthesis stages.

**Figure S3:** Variability of biomass and yield (grammes) between plant positions (P1-5) in the greenhouse for the nine barley varieties; Barque73, Buloke, Commander, Fathom, Flagship, Hindmarsh, Maritime, Schooner and Yarra.

**Figure S4:** Correlation between epigenetic distance using *Hpa*II (a, b) or *Msp*I (c, d) profiles and yield from control (a, c) and stress (b, d) plants (varieties: Barque73, Buloke, Commander, Fathom, Flagship, Hindmarsh, Maritime, Schooner and Yarra).

## Notes

### Competing Interest Statement

The authors have declared no competing interest.

