## Supplemental Figures for "Greenhouse spatial effects detected in the barley (*Hordeum vulgare* L.) epigenome underlie stochasticity of DNA methylation"

Supplementary Material

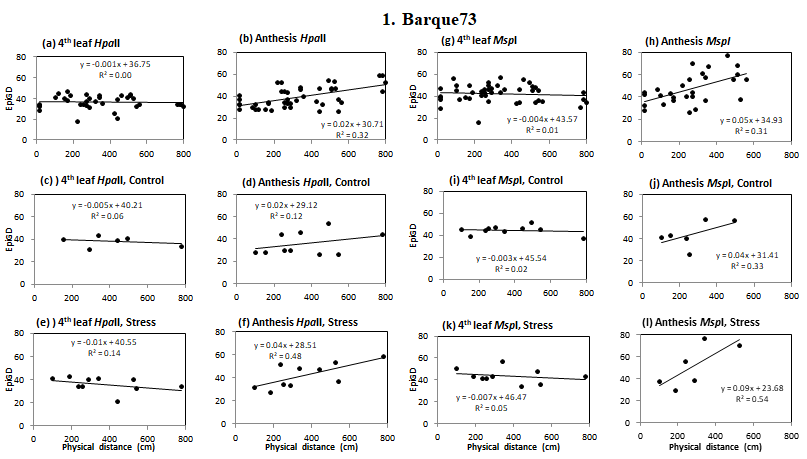

### Figure S1-1: Correlation between pairwise epigenetic distance (EpiGD) and physical distance between individual plants (cm, centimeter) of barley variety Barque 73, using the Mantel test and methylation sensitive enzymes *Hpa*II (a-f) and *Msp*I (g-l). Analyses involved control and stress plants together (a, b, g and h), control plants only (c, d, i and j) or stress plants only (e, f, k and l).

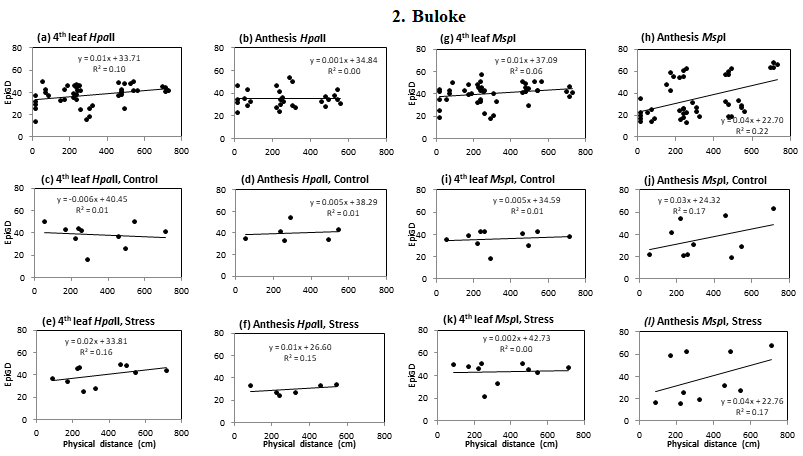

### Figure S1-2: Correlation between pairwise epigenetic distance (EpiGD) and physical distance between individual plants (cm, centimeter) of barley variety Buloke, using the Mantel test and methylation sensitive enzymes *Hpa*II (a-f) and *Msp*I (g-l). Analyses involved control and stress plants together (a, b, g and h), control plants only (c, d, i and j) or stress plants only (e, f, k and l).

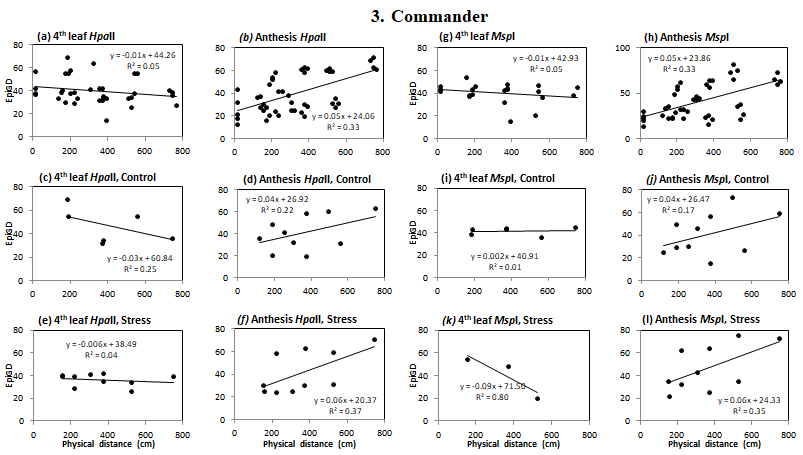

### Figure S1-3: Correlation between pairwise epigenetic distance (EpiGD) and physical distance between individual plants (cm, centimeter) of barley variety Commander, using the Mantel test and methylation sensitive enzymes *Hpa*II (a-f) and *Msp*I (g-l). Analyses involved control and stress plants together (a, b, g and h), control plants only (c, d, i and j) or stress plants only (e, f, k and l).

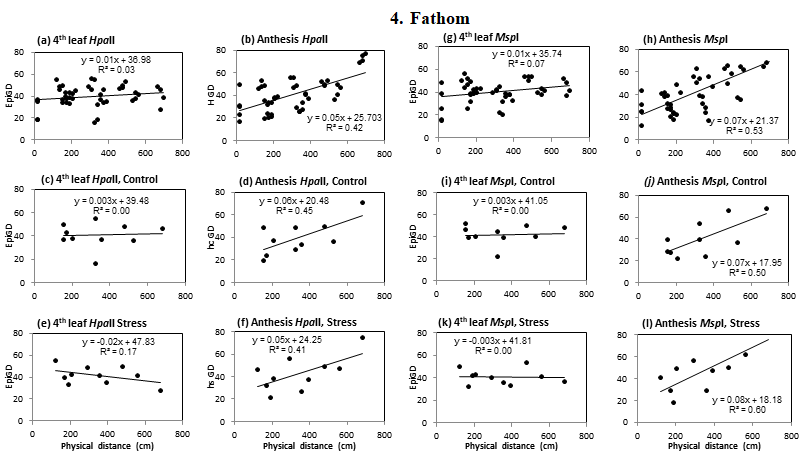

### Figure S1-4: Correlation between pairwise epigenetic distance (EpiGD) and physical distance between individual plants (cm, centimeter) of barley variety Fathom, using the Mantel test and methylation sensitive enzymes *Hpa*II (a-f) and *Msp*I (g-l). Analyses involved control and stress plants together (a, b, g and h), control plants only (c, d, i and j) or stress plants only (e, f, k and l).

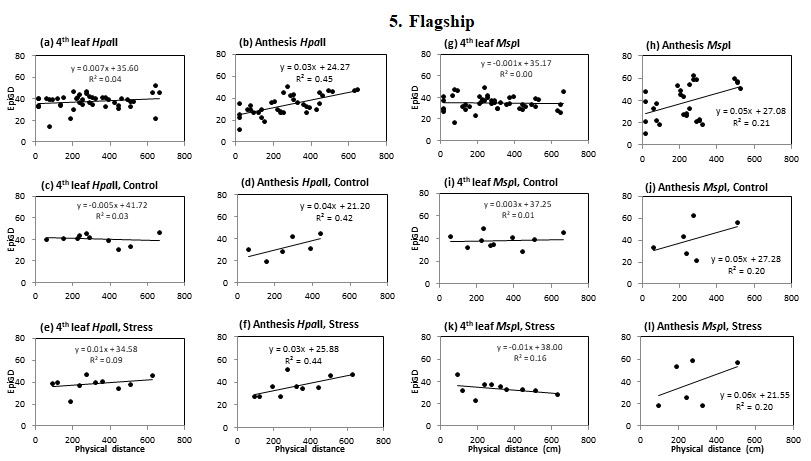

### Figure S1-5: Correlation between pairwise epigenetic distance (EpiGD) and physical distance between individual plants (cm, centimeter) of barley variety Flagship, using the Mantel test and methylation sensitive enzymes *Hpa*II (a-f) and *Msp*I (g-l). Analyses involved control and stress plants together (a, b, g and h), control plants only (c, d, i and j) or stress plants only (e, f, k and l).

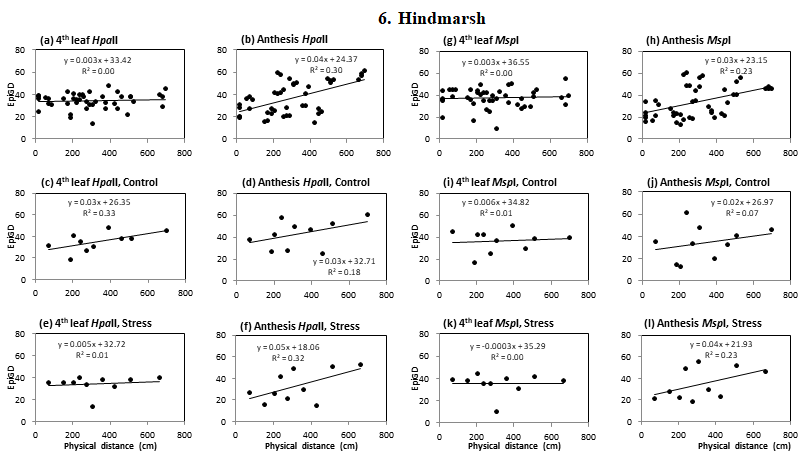

### Figure S1-6: Correlation between pairwise epigenetic distance (EpiGD) and physical distance between individual plants (cm, centimeter) of barley variety Hindmarsh, using the Mantel test and methylation sensitive enzymes *Hpa*II (a-f) and *Msp*I (g-l). Analyses involved control and stress plants together (a, b, g and h), control plants only (c, d, i and j) or stress plants only (e, f, k and l).

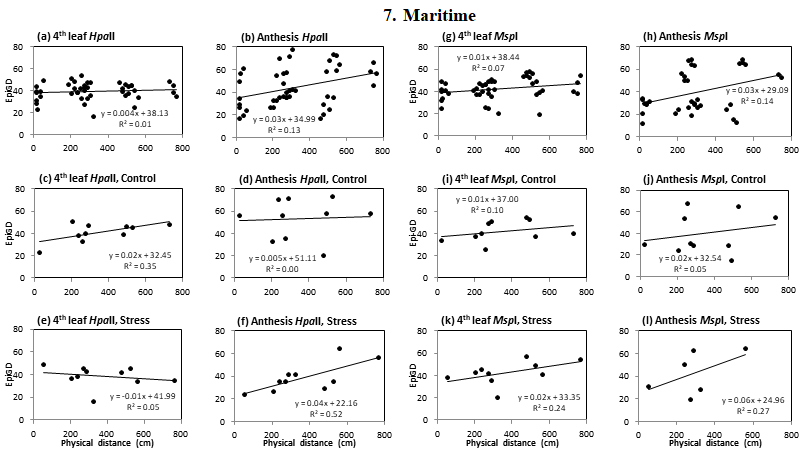

### Figure S1-7: Correlation between pairwise epigenetic distance (EpiGD) and physical distance between individual plants (cm, centimeter) of barley variety Maritime, using the Mantel test and methylation sensitive enzymes *Hpa*II (a-f) and *Msp*I (g-l). Analyses involved control and stress plants together (a, b, g and h), control plants only (c, d, i and j) or stress plants only (e, f, k and l).

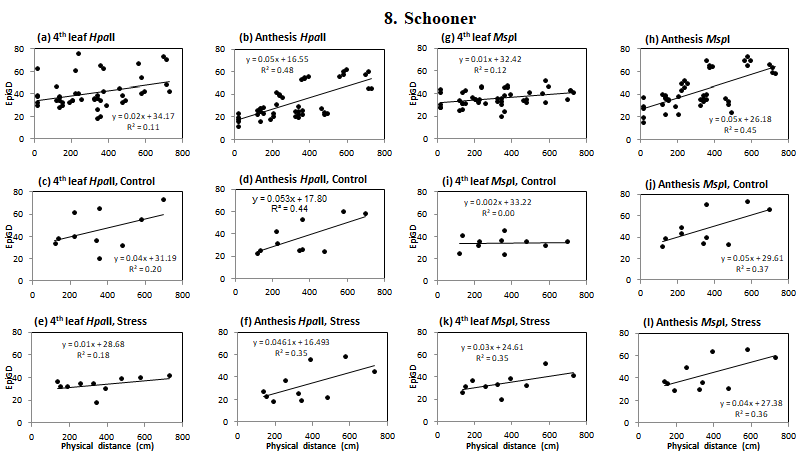

### Figure S1-8: Correlation between pairwise epigenetic distance (EpiGD) and physical distance between individual plants (cm, centimeter) of barley variety Schooner, using the Mantel test and methylation sensitive enzymes *Hpa*II (a-f) and *Msp*I (g-l). Analyses involved control and stress plants together (a, b, g and h), control plants only (c, d, i and j) or stress plants only (e, f, k and l).

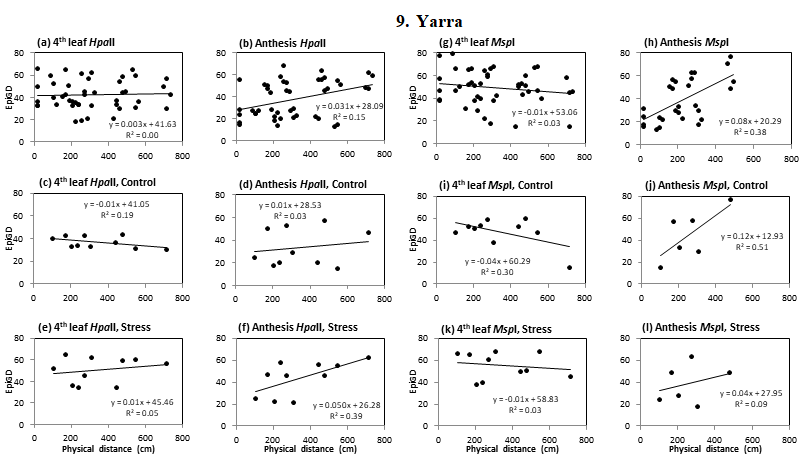

### Figure S1-9: Correlation between pairwise epigenetic distance (EpiGD) and physical distance between individual plants (cm, centimeter) of barley variety Yara, using the Mantel test and methylation sensitive enzymes *Hpa*II (a-f) and *Msp*I (g-l). Analyses involved control and stress plants together (a, b, g and h), control plants only (c, d, i and j) or stress plants only (e, f, k and l).

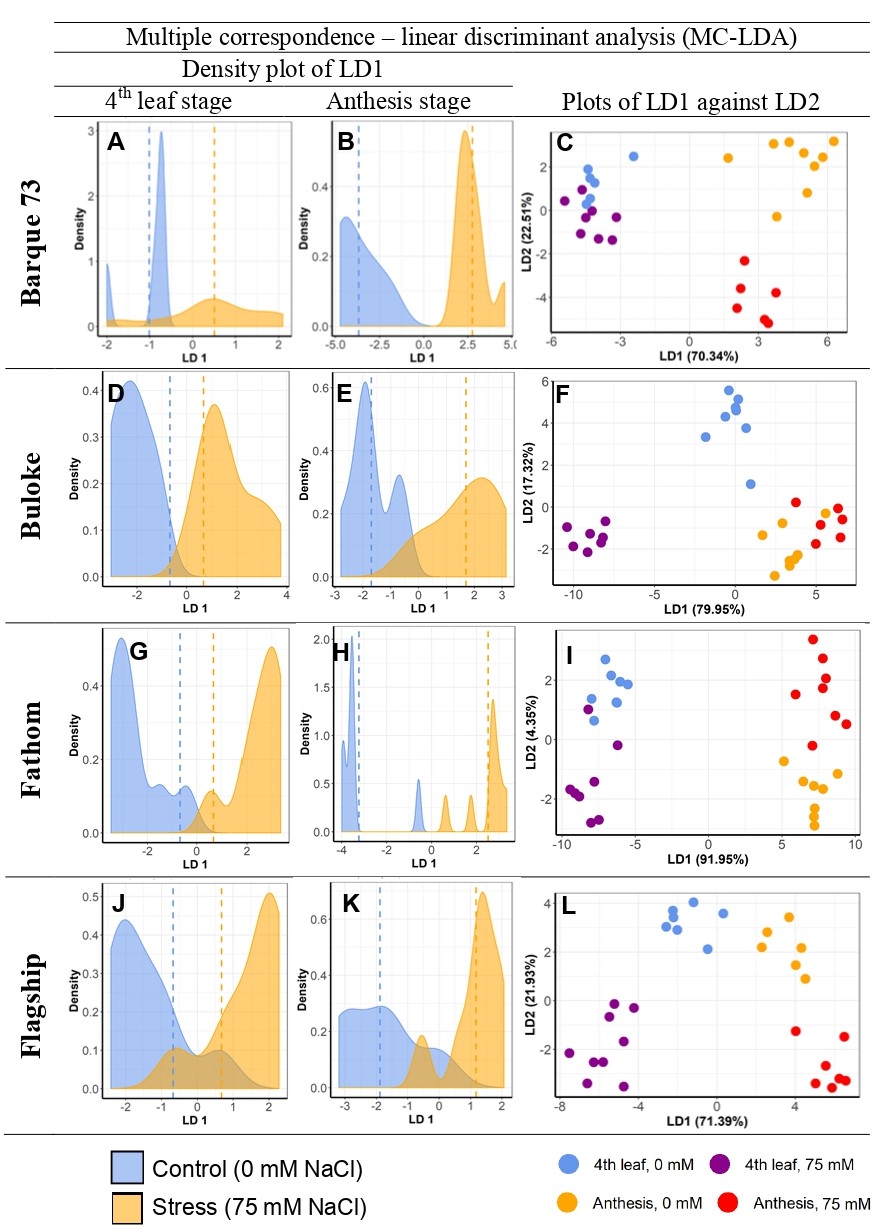

Figure S2: Multiple Correspondence and Linear Discriminant Analyses (MC-LDA) of MSAP markers in barley varieties (Barque73, Buloke, Fathom and Flagship) under salt stress (75 mM) and control (0 mM) conditions. The panel shows density plots of LD function between stress and control plants, at 4th leaf stage (A, D, G, J) and at anthesis (B, E, H, K). Dashed vertical lines represent the mean LD1 in 2 groups’ comparisons. Graphs of panel C, F, I, and L are MC-LDA plots comparing the salt treatment groups at both 4th leaf and anthesis stages.

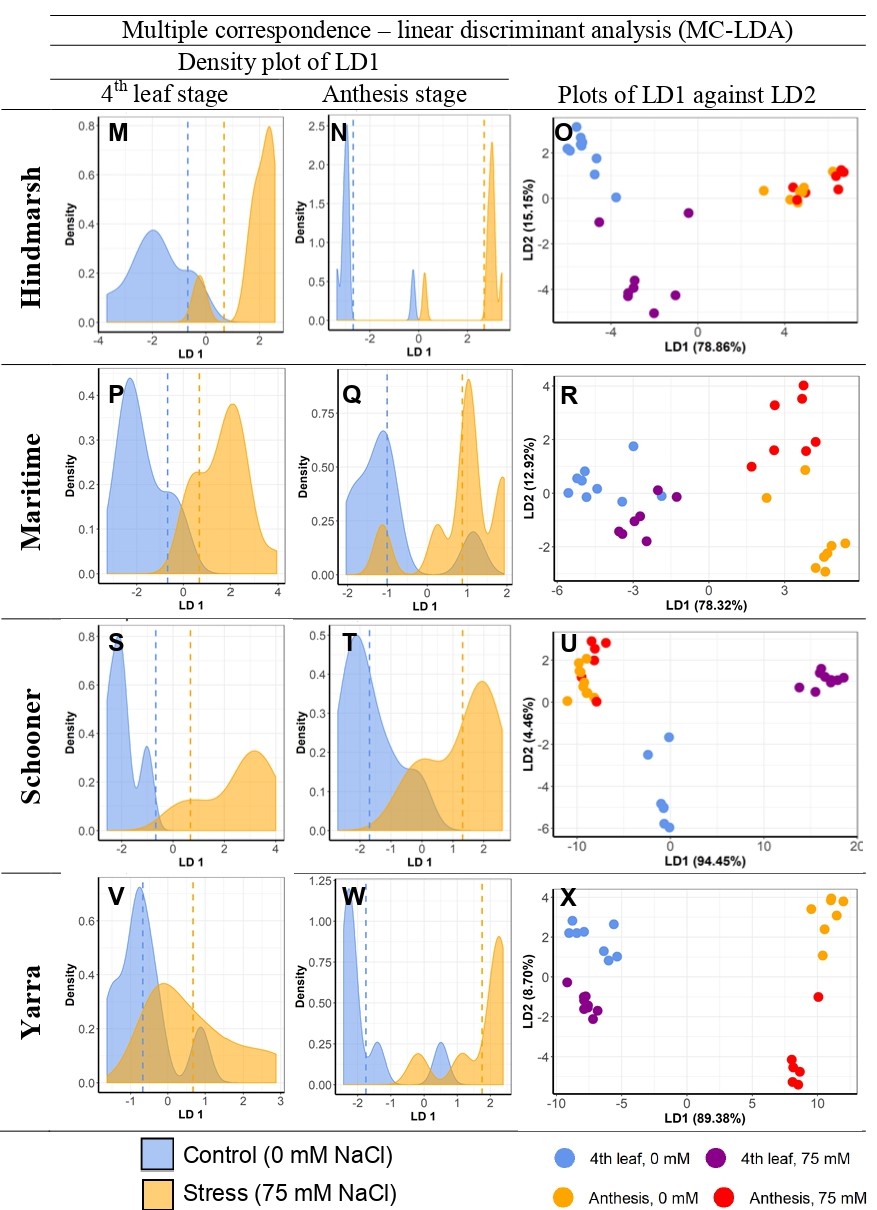

### Figure S2 (contd): Multiple Correspondence and Linear Discriminant Analyses (MC-LDA) of MSAP markers in barley varieties (Hindmarsh, Maritime, Schooner and Yarra) under salt stress (75 mM) and control (0 mM) conditions. The panel shows density plots of LD function between stress and control plants, at 4th leaf stage (M, P, S, V) and at anthesis (N, Q, T, W). Dashed vertical lines represent the mean LD1 in 2 groups’ comparisons. Graphs of panel O, R, U and X are MC-LDA plots comparing the salt treatment groups at both 4th leaf and anthesis stages.

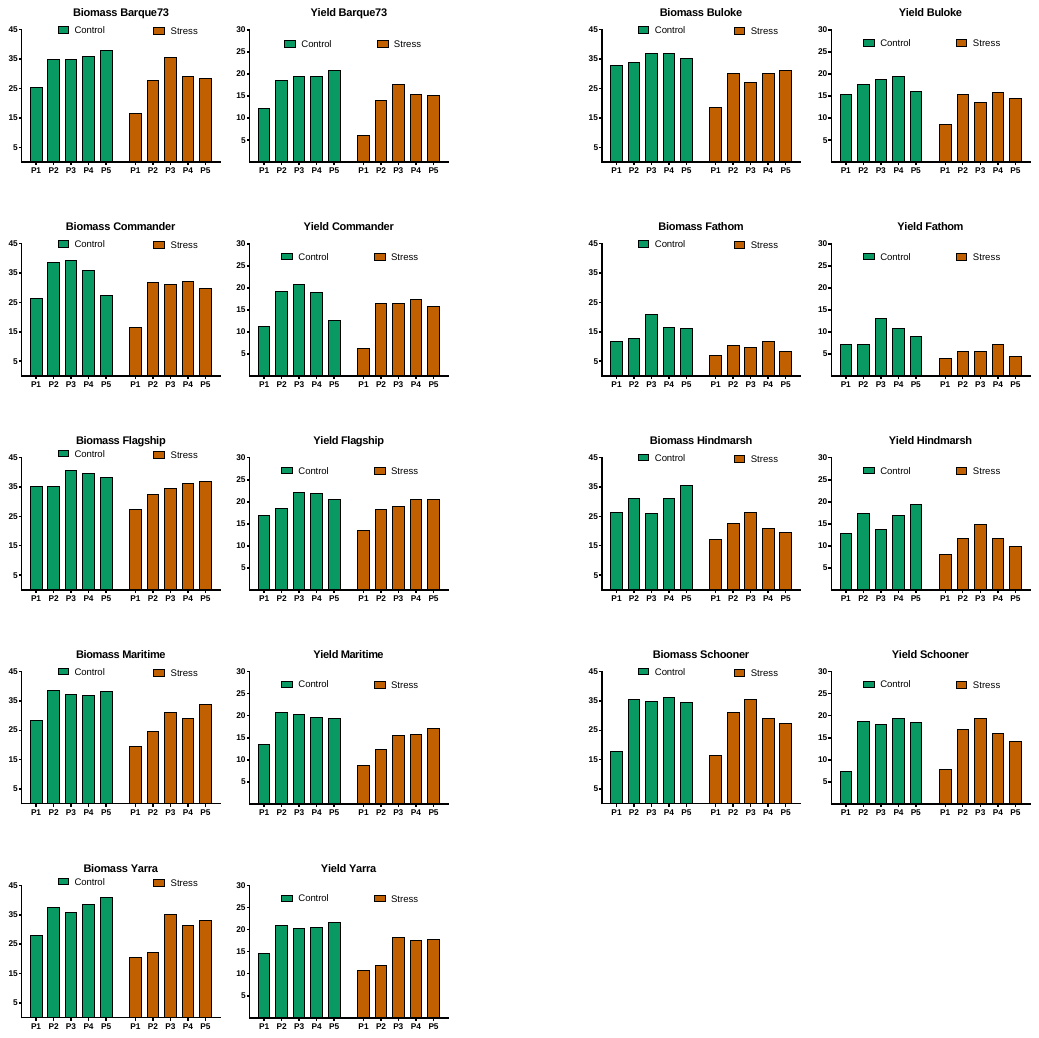

### Figure S3: Variability of biomass and yield (grammes) between plant positions (P1-5) of “control” and “stress” plants in the greenhouse for the nine barley varieties; Barque73, Buloke, Commander, Fathom, Flagship, Hindmarsh, Maritime, Schooner and Yarra.

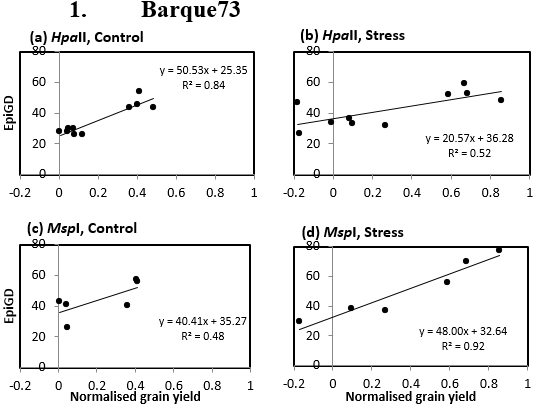

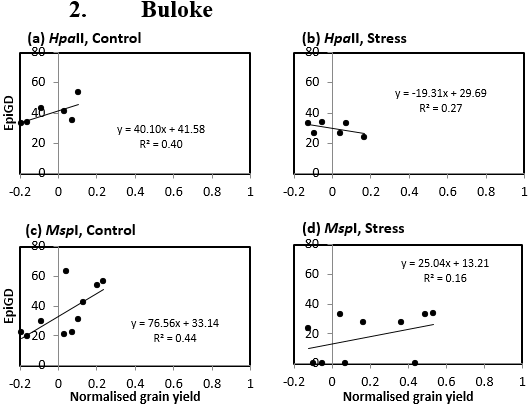

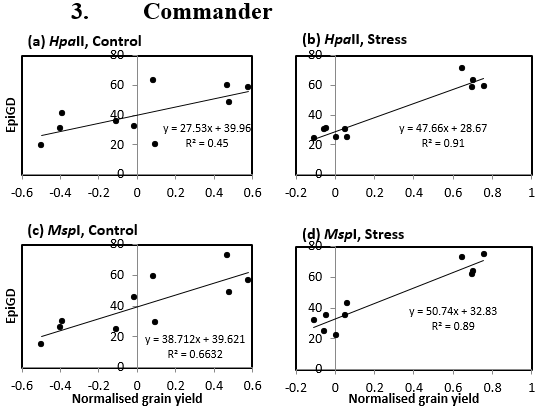

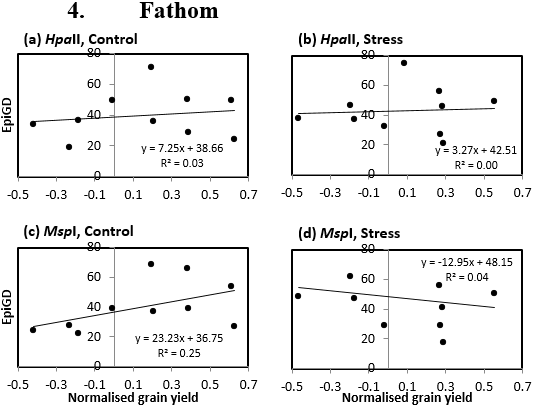

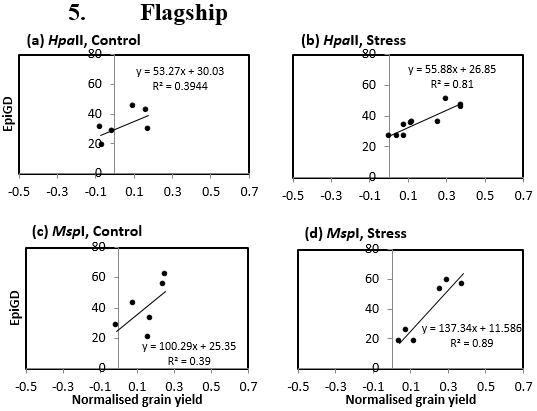

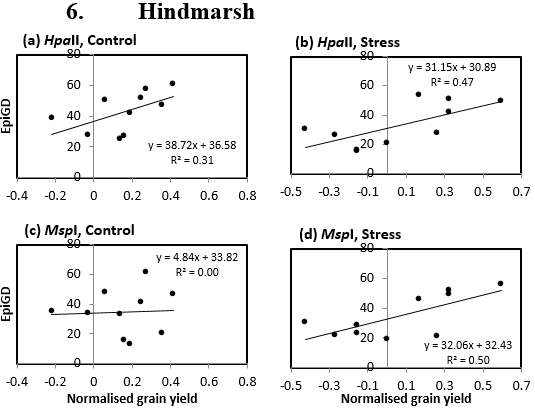

### Figure S4: Correlation between pairwise epigenetic distance using *Hpa*II (a, b) or *Msp*I (c, d) profiles and yield from control (a, c) and stress (b, d) plants (varieties; Barque73, Buloke, Commander, Fathom, Flagship, and Hindmarsh).

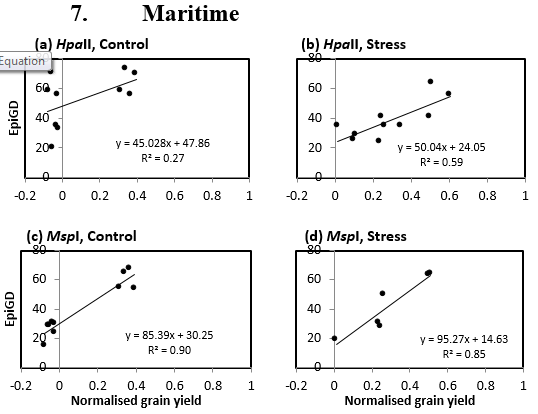

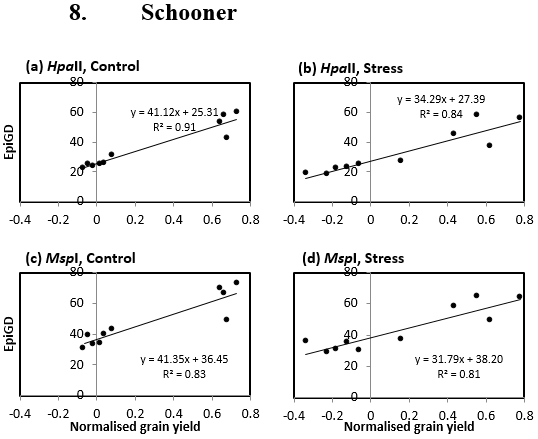

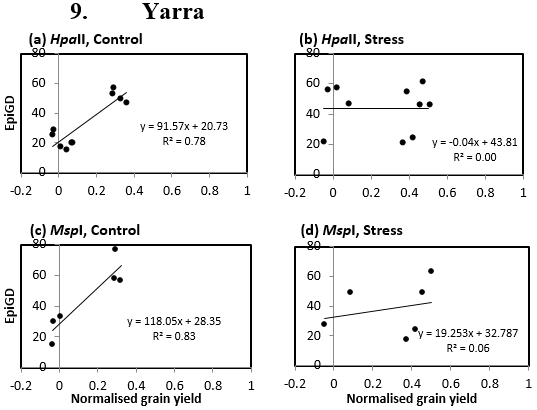

### Figure S4 (contd): Correlation pairwise between epigenetic distance using *Hpa*II (a, b) or *Msp*I (c, d) profiles and yield from control (a, c) and stress (b, d) plants (varieties Maritime, Schooner and Yarra).
